# Multimodal Connectome Markers of Disruptive Behavior Problems in Youth: Distinct Patterns of Concurrent Severity and Prospective Symptom Change

**DOI:** 10.64898/2026.10.26.761676

**Authors:** Eleni Christofilea, Kaiting Mai, Qinghao Liang, Raimundo X. Rodriguez, Caroline Lloyd, Yize Zhao, Dustin Scheinost, Denis G. Sukhodolsky, Karim Ibrahim

**Affiliations:** Child Study Center, Yale School of Medicine, New Haven, Connecticut, USA; Department of Biomedical Engineering, Yale University, New Haven, Connecticut, USA; Interdepartmental Neuroscience Program, Yale University, New Haven, Connecticut, USA; Department of Biostatistics, Yale School of Public Health, New Haven, Connecticut, USA; Department of Biomedical Informatics and Data Science, Yale School of Medicine, New Haven, Connecticut, USA; Wu Tsai Institute, Yale University, New Haven, Connecticut, USA; Yale Biomedical Imaging Institute, Yale School of Medicine, New Haven, Connecticut, USA; Department of Radiology and Biomedical Imaging, Yale School of Medicine, New Haven, Connecticut, USA; Department of Statistics and Data Science, Yale University, New Haven, Connecticut, USA; Department of Psychology, Yale University, New Haven, Connecticut, USA; Department of Social and Behavioral Sciences, Yale School of Public Health, New Haven, Connecticut, USA

## Abstract

Disruptive behaviors problems, such as irritability/anger and maladaptive aggression, are associated with altered connectivity across networks supporting emotion regulation. However, multimodal connectome markers of concurrent severity and prospective symptom change, including sex-specific patterns, remain unclear. Using a large, sex-balanced sample from the Adolescent Brain Cognitive Development (ABCD) Study, we applied multimodal connectome-based predictive modelling (CPM) to test whether resting-state functional (*n* = 7,556; 49.3% female) and structural (*N* = 8,978; 48.4%) connectivity predicted baseline disruptive behavior severity (at ages 9-10) and 2-year symptom change (from ages 9-10 to 11-12). Models were adjusted for covariates, permutation-tested, and FDR corrected. Secondary analyses tested sex-stratified models separately in males and females. In models emerging as significant, follow-up analyses tested for distinct vs shared predictive edges. Functional and structural connectomes predicted baseline severity (*ρ* = 0.087 and *ρ* = 0.071, respectively). Functional connectivity also predicted prospective symptom change (*ρ* = 0.038), whereas prospective structural prediction was predominately with negative edges (*ρ* = 0.026). Functional connectivity models were significant in both sexes, while prospective structural connectivity models were significant in males, but not females. Frontoparietal, medial frontal, sensorimotor, salience, cerebellar, and visual networks contributed across models. Cross-sectional models showed a more distributed pattern while prospective models showed a more focal pattern. These findings identify complementary, largely distinct functional and structural connectome patterns associated with current disruptive behavior risk and subsequent symptom change, supporting the importance of developmental timing, sex, and imaging modality in developing brain-based biomarkers of disruptive behavior risk in youth.

## Introduction

Disruptive behavior disorders, characterized by irritability/anger, aggression, noncompliance, and/or antisocial behavior [1], are a prevalent psychiatric condition in youth [2,3], carrying a profound public health impact and societal costs [4-7]. Youth with disruptive behavior are at a higher risk of adverse outcomes including substance abuse and risk of other co-occurring psychiatric conditions, contributing to significant impairment and distress and accounting for the most common reason for clinical referrals [6,8,9]. Disruptive behavior problems also constitute a transdiagnostic dimension [10,11] that co-occurs across several child psychiatric disorders [12,13]. Despite this clinical importance, it remains unclear whether functional and structural connectomes provide shared or complementary markers of concurrent disruptive behavior severity and subsequent symptom change.

### Functional and Structural Circuit Markers of Disruptive Behavior in Youth

Disruptive behavior problems are associated with difficulties in emotion regulation in youth [14]. On a neural-systems level, disruptive behavior has been consistently linked to perturbations in functional connectivity across networks supporting emotion regulation and broader cognitive control processes, particularly frontoparietal and frontolimbic circuitry [15-24]. In a clinical sample of youth, we showed that reduced connectivity in frontoparietal nodes spanning the dorsal and ventral prefrontal cortex and posterior parietal cortex is linked to greater disruptive behavior symptoms in children and adolescents during emotional face processing [17,18]. These findings have been corroborated by connectome-based studies, which further suggest that disruptive behavior severity is associated with distributed broader network dysfunction spanning frontoparietal, default mode, sensorimotor, and salience networks, with frontoparietal and related dorsal and ventral prefrontal nodes emerging as prominent predictive and generalizable features [19].

Similar findings have been shown with resting-state functional MRI (fMRI), with aberrant connectivity reported across frontoparietal, default mode, salience, limbic, and social cognitive networks in youth with elevated disruptive behavior symptoms [23,25-32]. Adding to these findings, recent work from our group showed that alterations in the dynamic, time-varying properties of resting-state functional connectivity were related to symptom severity, whereby children with elevated disruptive behavior showed greater occupancy time within globally dysconnected brain states, marked by aberrant connectivity, particularly within frontoparietal circuitry, and reduced frontoparietal integration with other networks [33]. Work from the few existing longitudinal imaging studies of disruptive behavior suggest that altered connectivity across frontolimbic, cingulo-opercular, and salience nodes is associated with worsening disruptive behavior over time [31,34]. However, most prior work with functional connectivity has been cross-sectional and/or focused on predefined regions of interest. Consequently, it remains unknown whether baseline brain-wide connectivity (or connectomes) predict subsequent changes in disruptive behavior, and whether patterns of functional connectivity associated with concurrent severity differ from patterns associated with prospective symptom change.

Similarly, microstructural alterations in the integrity of white matter tracts are also related to disruptive behavior in youth, predominately in association fibers across frontoparietal and frontolimbic nodes supporting emotion regulation and broader cognitive control spanning prefrontal, parietal, temporal, and limbic regions [35-42]. For instance, increased disruptive behavior severity in youth is associated with reduced microstructural integrity of the longitudinal fasciculi [35,37,39,43], cingulum [35,37,39,44,45], and uncinate fasciculus [35,36,38,39,43,46]. Our group found that youth with maladaptive aggression show altered microstructural integrity across association tracts, including the cingulum, relative to unaffected controls [45]. Aligned with diffusion MRI studies, findings from recent mega-analyses showed neuroanatomical structural alterations in frontoparietal, salience, and default mode circuitry in children and adolescents with elevated disruptive behavior [47-49]. While neuroanatomical studies show alterations frontoparietal, frontotemporal, and frontolimbic nodes linked to worsening disruptive behavior [37,50-53], the few longitudinal diffusion MRI studies further suggest that perturbations in the organization and integrity of frontoparietal and limbic nodes is associated with subsequent worsening of disruptive behavior [37,54,55]. While findings converge on a plausible dysfunction in microstructural integrity, particularly across frontoparietal nodes, prior diffusion MRI studies have predominantly examined selected tracts or regional measures rather than brain-wide structural connectomes. Thus, it remains unclear whether structural connectome organization predicts concurrent and prospective disruptive behavior or provides information complementary to functional connectivity.

### Present Study

The present study used resting-state functional and diffusion MRI data from the Adolescent Brain Cognitive Development (ABCD) Study to identify multimodal connectome markers of disruptive behavior problems. First, data-driven, connectome-based predictive modeling (CPM) was used to test whether baseline functional and structural connectivity predicted concurrent disruptive behavior severity at ages 9-10 and predicted two-year change in severity from ages 9-10 to 11-12. Next, exploratory sex-stratified analyses were used to examine predictive connectome patterns separately in males and females. Prior neuroimaging studies of disruptive behavior have often included predominantly male samples and/or lacked sufficient statistical power to characterize connectivity patterns separately in males and females [15,56]. Given the mixed findings regarding sex-related variation in functional and structural markers of disruptive behavior [18,21,28,36,47-49,55,57-63], sex-stratified analyses were exploratory (see **Supplemental Background**). Informed by prior work [19], we predicted that functional and structural connectivity would predict concurrent severity and prospective symptom change, with predictive connections distributed across networks supporting emotion regulation, cognitive control broadly, and social perception spanning frontoparietal, medial frontal, salience, default mode, and sensorimotor circuits. We also expected that concurrent and prospective models would show partially distinct connectome configurations, with frontoparietal circuity as highly predictive. While sex-stratified analyses were exploratory, based on prior work [18,28,36,61], we expected sex-specific multimodal predictive patterns for cross-sectional and prospective models.

## Materials and Methods

### Participants

The current study used baseline (9-10 years) and two-year follow-up (11-12 years) data from the ABCD Study® (Data Release 5.1), a longitudinal study of youth recruited across 21 sites in the United States [64-67]. Participants were required to have complete behavioral data and resting-state functional and/or diffusion-weighted imaging data meeting modality-specific and quality control criteria. Of 11,878 enrolled participants, 10,988 had available resting-state fMRI and/or diffusion-weighted imaging data. Consistent with prior ABCD connectome studies [19,33], functional imaging analyses were restricted to participants with mean framewise displacement ≤0.25 mm and valid connectome data. The final samples included 7,556 participants with functional connectomes (3,726 females) and 8,978 participants with structural connectomes (4,343 females). Participant characteristics are presented in **Table 1**. Additional sample selection details are provided in the **Supplemental Methods**.

**Table 1.**
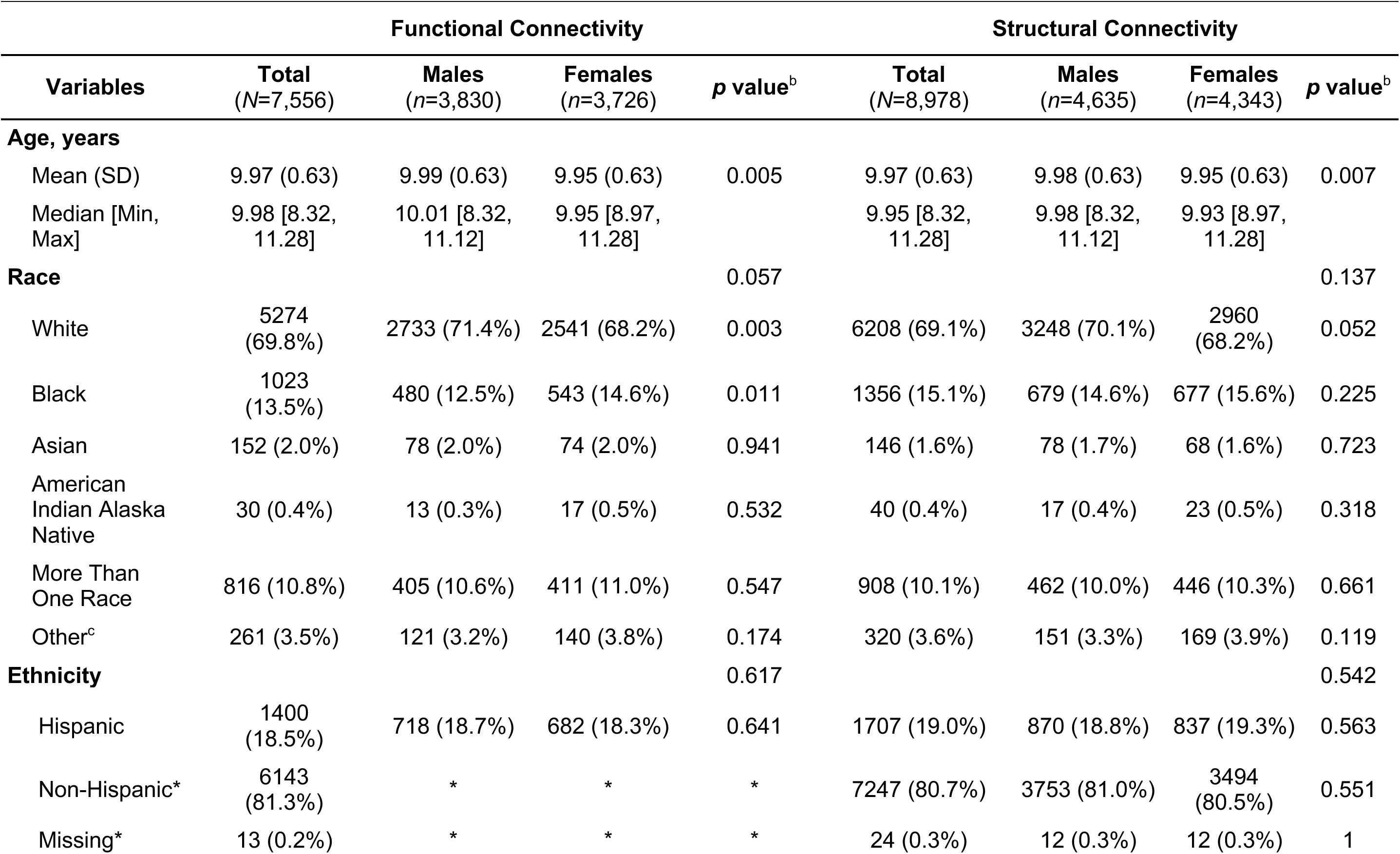
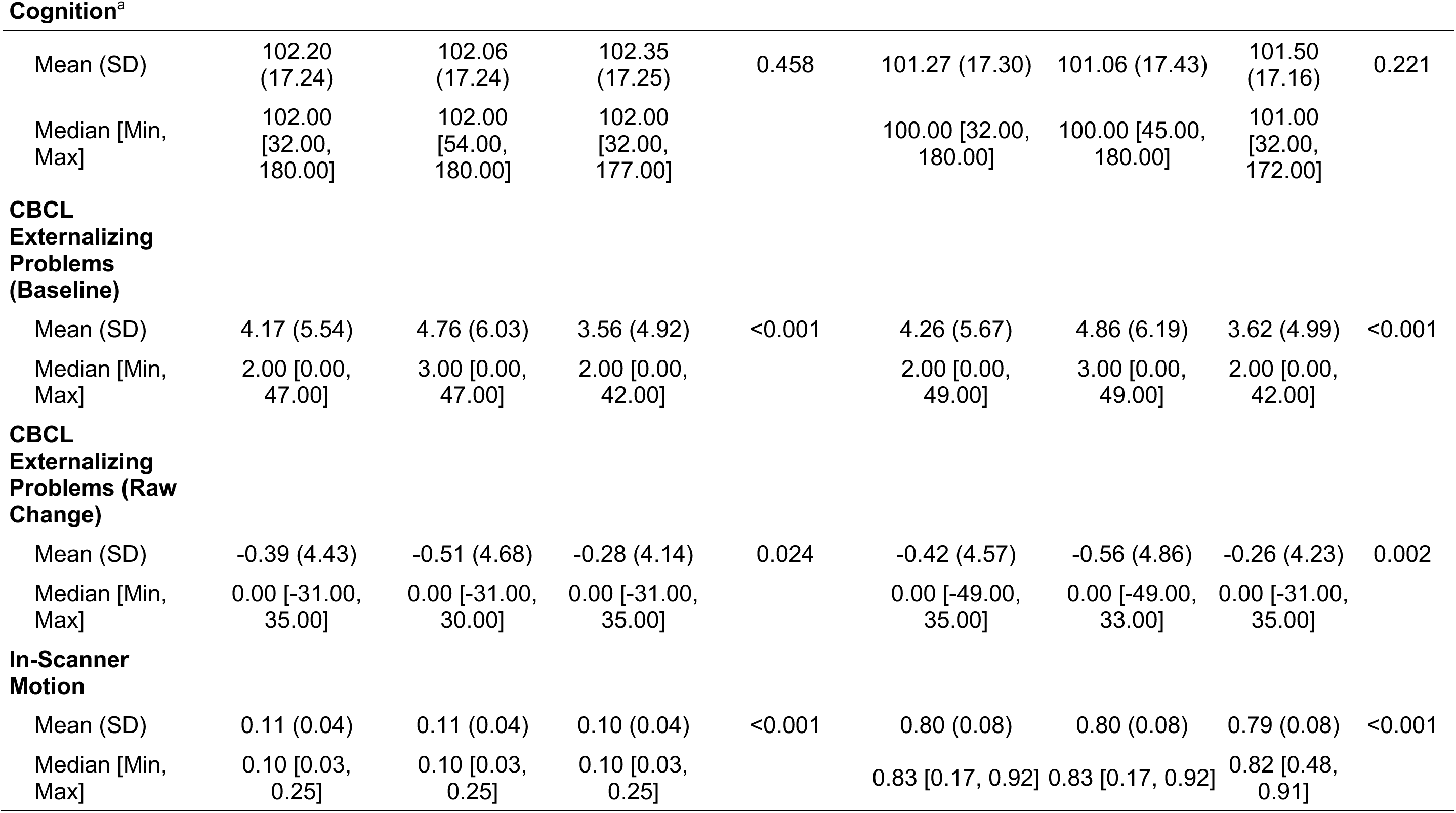
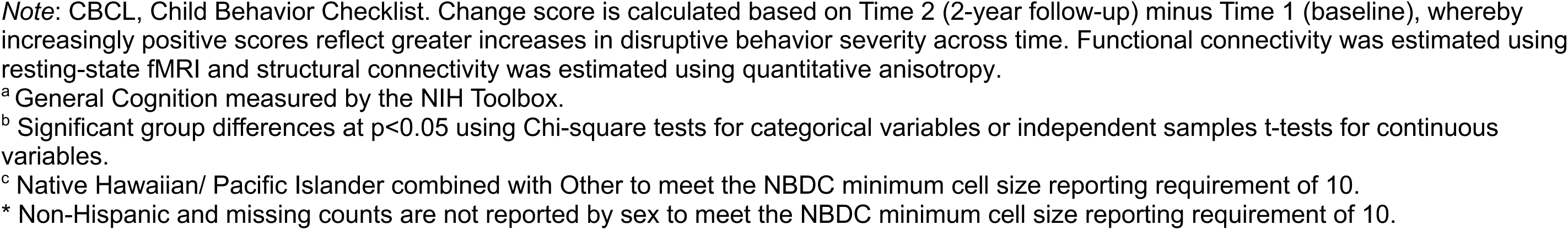
Participant Demographics and Clinical Characteristics.

### Behavioral Measures

The parent-rated Child Behavior Checklist (CBCL) Externalizing Behavior Problems subscale [68] was used as a continuous measure of disruptive behavior severity, consistent with previous ABCD neuroimaging studies [19,21,33,69,70]. The NIH Toolbox Cognition battery total composite score [71] was included in analyses to account for cognition.

Additional measure details are provided in the **Supplemental Methods**.

### Neuroimaging Data Acquisition and Preprocessing

Neuroimaging data using structural T1-weighted scan, resting-state, and diffusion MRI were obtained from the ABCD Study [64,65]. Participants completed at least 8 minutes of eyes open resting-state fMRI and diffusion-weighted imaging was acquired using the ABCD multishell diffusion protocol. For details on the ABCD imaging protocol, see Hagler et al. [65] and **Supplemental Methods**.

Functional and structural connectomes were generated using the Shen 268-node atlas [72], yielding one 268 × 268 connectivity matrix per participant and modality. Resting-state functional connectivity was defined as Fisher z-transformed pairwise correlations between regional BOLD time series after established preprocessing, nuisance regression, global signal regression, motion correction, and temporal filtering [19,73-76]. Structural connectivity was reconstructed from diffusion-weighted imaging using generalized q-sampling imaging and deterministic tractography and was quantified as mean quantitative anisotropy for streamlines connecting each regional pair [77,78]. The same parcellation was used across modalities, and modality-specific motion and image quality measures were included as covariates [70]. Full details for preprocessing and connectome construction procedures are provided in the **Supplemental Methods**.

### Analytic Approach

#### Connectome-based Predictive Modeling

Connectome-based predictive modeling (CPM) was conducted separately for baseline functional and structural connectivity to test prediction of concurrent disruptive behavior severity and two-year symptom change [72,74,75,79]. Models used 10-fold cross-validation repeated 100 times. Within each training fold, edges positively or negatively associated with the outcome after adjustment for model-specific covariates at *p* ≤ 0.01 were retained in separate feature masks, and feature selection was confined to the training data [72,79,80]. For each participant, positive and negative network strength scores were estimated by summing connectivity weights across edges in the corresponding feature masks. Combined network strength was calculated as positive minus negative network strength and entered as a single predictor in a robust linear regression model. Positive and negative edge models were also fitted separately using the corresponding network strength scores. Fitted models were applied to held-out participants to generate predicted behavioral scores. Predictive performance was quantified using Spearman correlations between observed and predicted scores, with performance from a median ranked cross-validation repetition retained for reporting. Statistical significance was evaluated using 1,000 permutations of the behavioral labels, and permutation-derived *p* values were adjusted for multiple comparisons using Benjamini-Hochberg false discovery rate correction. Additional details are provided in the **Supplemental Methods**.

### Disruptive Behavior Connectome Models

Concurrent models predicted baseline CBCL Externalizing Problems raw scores as the dependent variable. Prospective models predicted symptom change as the dependent variable, calculated as the two-year follow-up score minus the baseline score, with higher values indicating greater symptom increases or less symptom reduction. Baseline functional and structural connectomes were analyzed separately in the total sample and in exploratory sex-stratified models. Models adjusted for age, cognition, and modality-specific motion. Regarding accounting for head motion in models, functional models used mean framewise displacement [81] and structural models used masked neighboring diffusion-weighted imaging correlation [82]. Predictive models in the total sample adjusted for biological sex, and prospective models additionally adjusted for baseline disruptive behavior severity (see also **Supplemental Methods**).

Predictive networks from significant CPM models were summarized at the edge, node, and canonical subnetwork levels [19,76]. Edge-level summaries identified individual connections selected by CPM. Node-level summaries quantified the number of predictive edges incident on each Shen atlas node. Subnetwork-level summaries were generated by assigning Shen atlas nodes to 10 canonical subnetworks and estimating the distribution of selected edges within and between subnetworks, following established approaches for connectome interpretation [76,83,84]. Overlap of nodes with macroscale brain regions (e.g., prefrontal cortex) was based on established anatomical labels [73] and overlap of nodes with canonical network localizations (e.g., frontoparietal) was based on the networks delineated in our prior work [85].

### Subnetwork-Level Comparison of CPM Feature Masks

To assess the extent to which CPM-derived predictive networks showed shared vs model-specific topographic organization, feature masks were compared at the subnetwork level. The follow-up contrasts included functional and structural connectivity models (edges) that emerged as significant in predicting baseline disruptive behavior and prospective symptom change. Benjamini-Hochberg false discovery rate correction was applied. Parcellation and statistical significance testing procedures are detailed in the **Supplemental Methods.**

## Results

### Cross-Sectional Models

#### Prediction results

##### Functional connectivity

Baseline functional connectivity predicted concurrent disruptive behavior severity in the total sample (*ρ* = 0.087, RMSE = 5.660, *p_FDR_* = 0.0035) (**Table S1**; **Figure 1**, top panel; **Figure 2**, left panel). In sex-stratified analyses, functional connectivity predicted concurrent severity in both male and female subsamples (**Figure 1**; **Figure 3**), with predictive performance numerically higher in males (*ρ* = 0.082) than females (*ρ* = 0.037) (**Table S1**).

**Figure 1.**
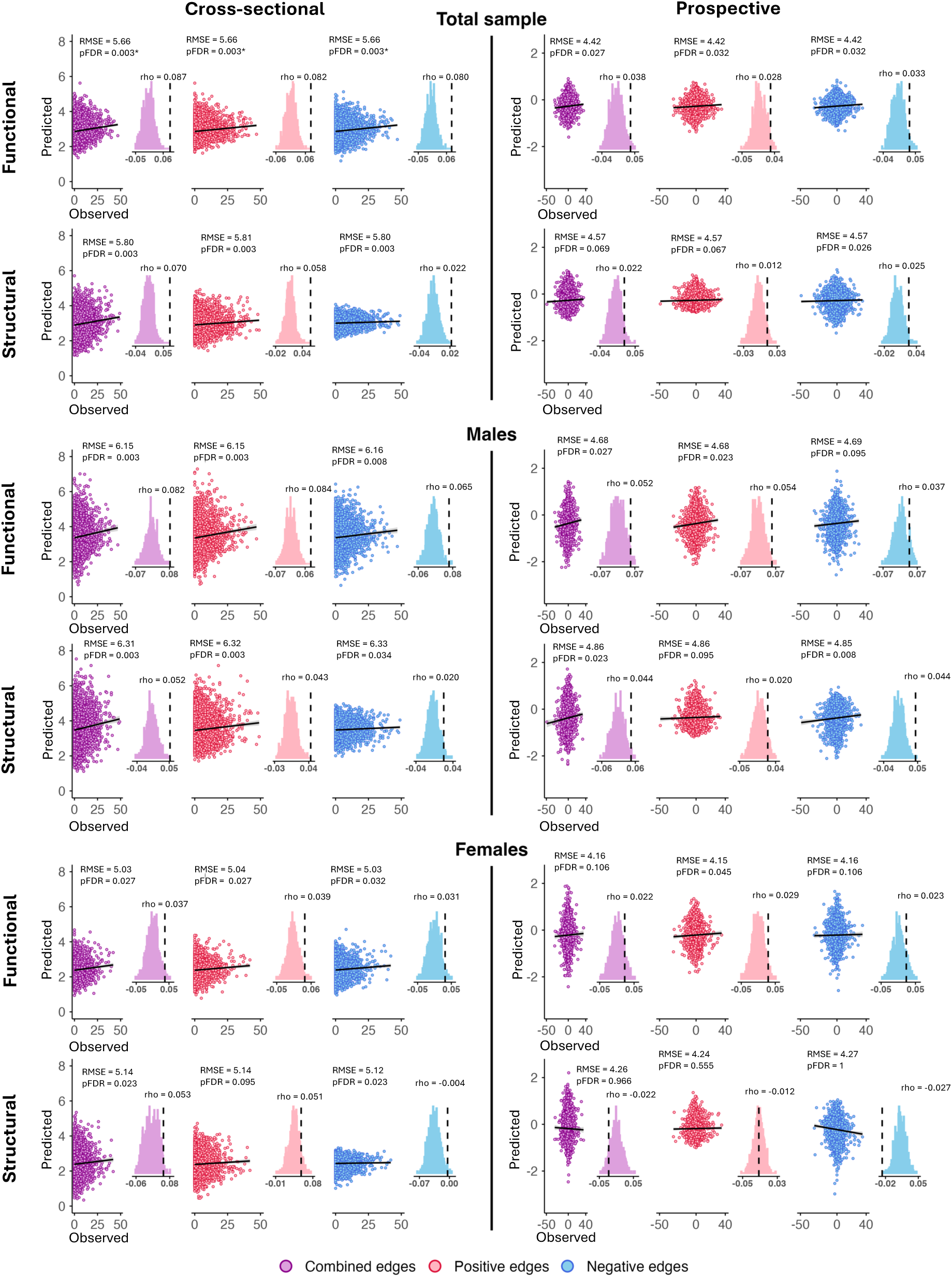
Multimodal connectome predicts disruptive behavior problems in youth. Cross-sectional models assessed prediction of disruptive behavior severity at baseline from functional and structural connectomes (left panel). Prospective models assessed prediction of change in disruptive behavior severity from baseline to the two-year follow-up from baseline connectomes using a raw change score (right panel). The Child Behavior Checklist (CBCL) Externalizing Behavior Problems subscale was used as a continuous measure of severity of disruptive behavior at baseline for cross-sectional models and at two-year follow-up for prospective models using a CBCL raw change score (i.e., two-year follow-up score minus baseline score). All models accounted for age, cognition, modality-specific motion, biological sex (in the total sample), and additionally for baseline disruptive behavior severity in prospective models. Scatterplots show observed (x-axis) vs predicted (y-axis) disruptive behavior. Histograms show the distribution of Spearman rho values from 1,000 iterations of randomly shuffled ratings of disruptive behavior severity used to nonparametrically determine *p* values (dashed line indicates the Spearman rho for the true model). Inset figures show the permutation distribution of Spearman rho values. The *p* values shown were obtained from permutation testing and then FDR corrected across tests within each modality (functional connectivity, structural connectivity). Each panel shows combined positive and negative edges (purple) as well as separate positive (red) and negative (blue) edges predicting disruptive behavior. For positive networks, increased edge weights (i.e., increased connectivity) predict greater severity of disruptive behavior. For negative networks, decreased edge weights (i.e., decreased connectivity) predict greater severity of disruptive behavior. Note: RMSE = Root Mean Square Error.

**Figure 2.**
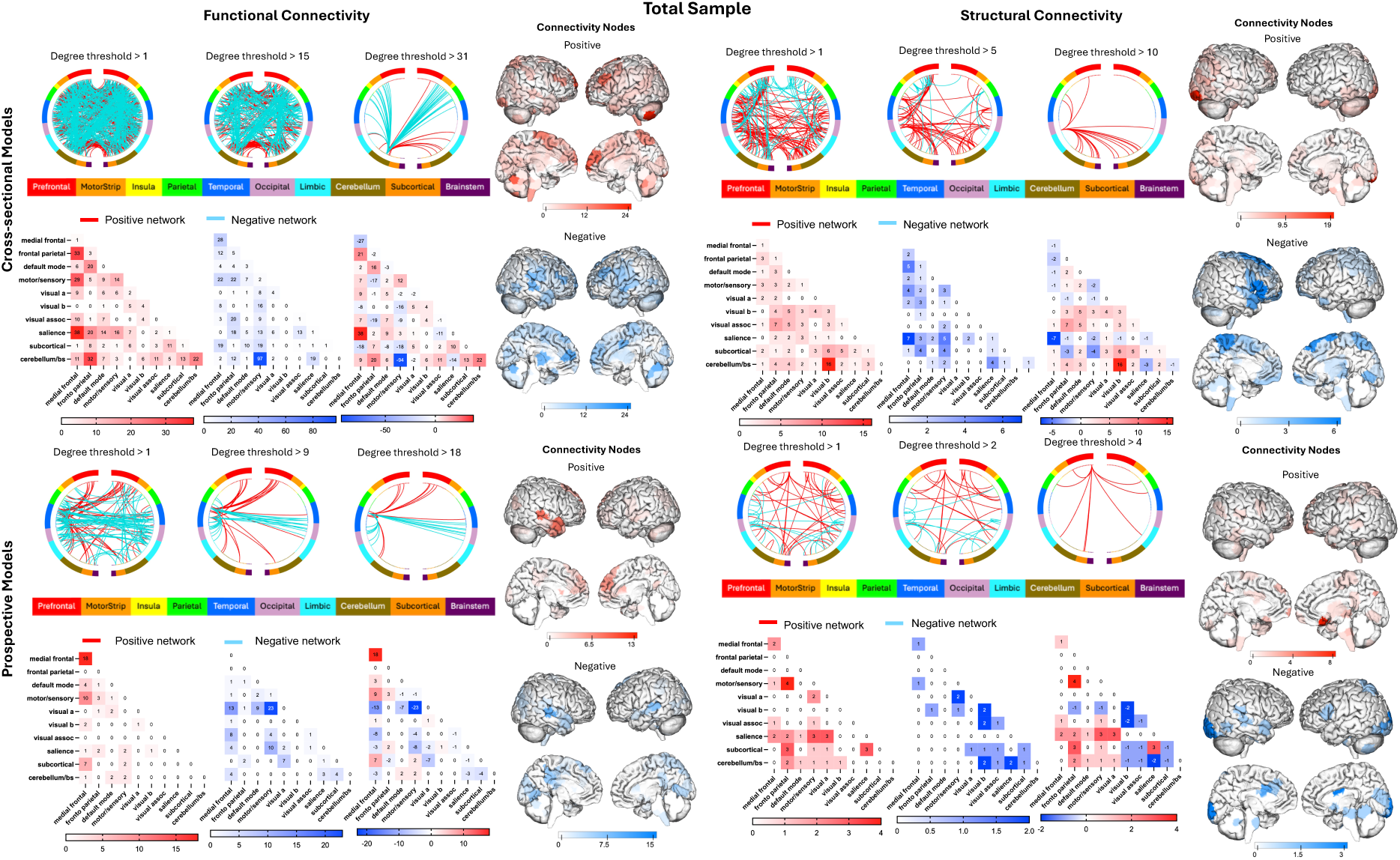
Networks predicting concurrent and prospective disruptive behavior problems in the total sample. Networks are summarized by functional and structural connectivity between macroscale brain regions, nodes, and networks. For circle plots, brain regions are presented in approximate anatomical order whereby longer-range connections are represented by longer lines. To visualize these complex networks while clarifying edges that contributed the most to the CPM model, varying thresholds are implemented in circle plots (e.g., edges belonging to nodes with 15 or more edges are denoted as degree threshold ≥15). For rendered brains, visualization of node degree (i.e., the sum of predictive edges for a node) for the positive and negative networks are shown. Darker colors indicate higher degree nodes (i.e., with more edges) contributing to the CPM model. For network matrices, within-and between-network connectivity for positive networks (red), negative networks (blue), and combined networks (depicts red and blue) are summarized using canonical networks. Cells represent the total number of edges connecting nodes within and between each network, with darker colors indicating a greater number of edges. By definition, positive and negative networks do not contain overlapping edges. Functional connectivity was estimated using resting-state fMRI and structural connectivity was estimated using quantitative anisotropy.

**Figure 3.**
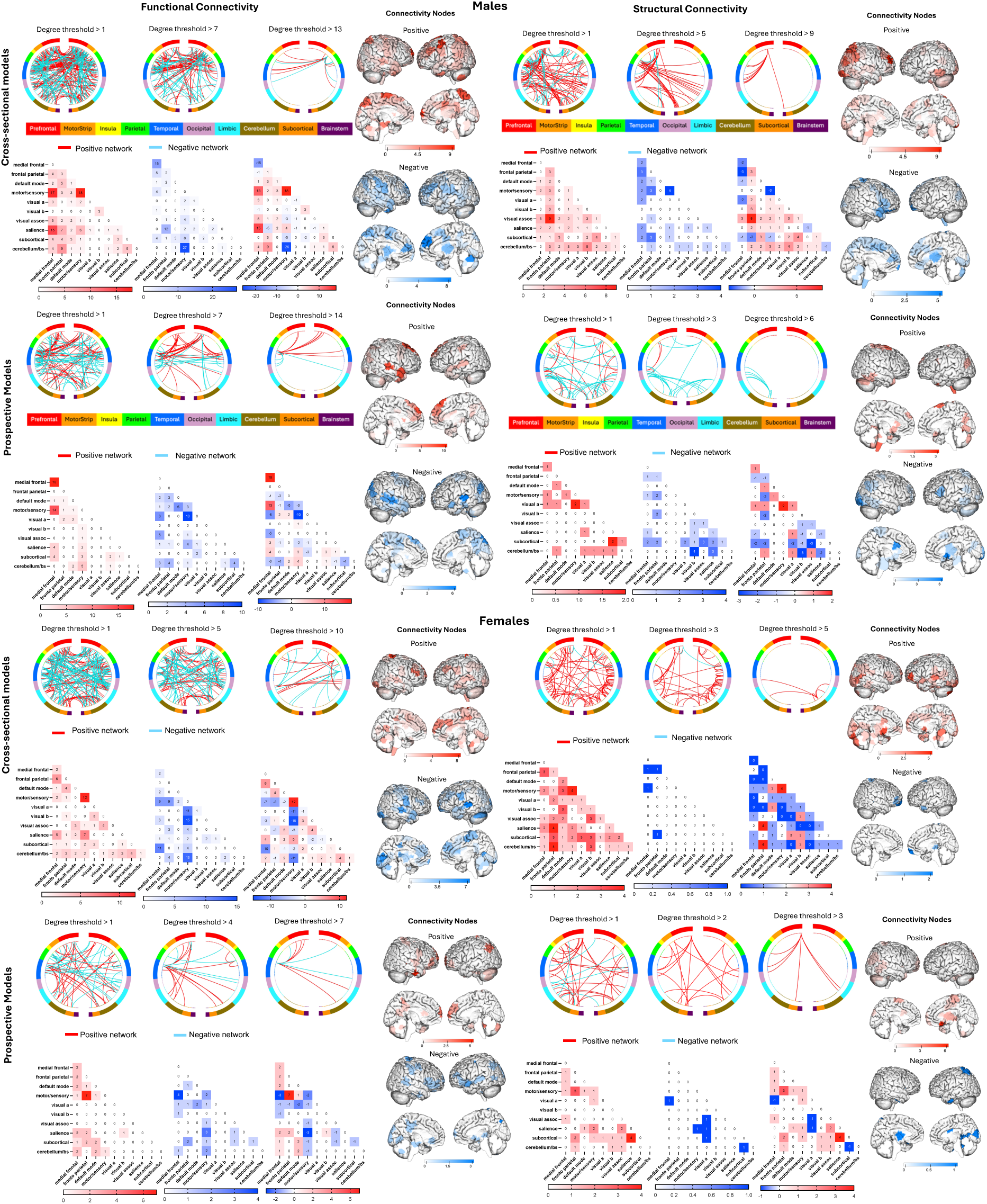
Sex-specific networks of concurrent and prospective disruptive behavior problems in the total sample. For male (top panel) and female (bottom panel) subgroups, networks are summarized by functional and structural connectivity between macroscale brain regions, nodes, and networks. For circle plots, brain regions are presented in approximate anatomical order whereby longer-range connections are represented by longer lines. To visualize these complex networks while clarifying edges that contributed the most to the CPM model, varying thresholds are implemented in circle plots (e.g., edges belonging to nodes with 15 or more edges are denoted as degree threshold ≥15). For rendered brains, visualization of node degree (i.e., the sum of predictive edges for a node) for the positive and negative networks are shown. Darker colors indicate higher degree nodes (i.e., with more edges) contributing to the CPM model. For network matrices, within-and between-network connectivity for positive networks (red), negative networks (blue), and combined networks (depicts red and blue) are summarized using canonical networks. Cells represent the total number of edges connecting nodes within and between each network, with darker colors indicating a greater number of edges. By definition, positive and negative networks do not contain overlapping edges. Functional connectivity was estimated using resting-state fMRI and structural connectivity was estimated using quantitative anisotropy.

##### Structural connectivity

Baseline structural connectivity predicted concurrent disruptive behavior severity (*ρ* = 0.071, RMSE = 5.801, *p_FDR_* = 0.0035), with positive edges contributing more (*ρ* = 0.058 for positive edges and *ρ* = 0.022 for negative edges) (**Table S1**; **Figure 1**, bottom panel). In sex-stratified analyses, structural connectivity predicted concurrent severity in both male and female subsamples (*ρ* = 0.05 for males and *ρ* = 0.05 for females) (**Figure 1**; **Figure 3**; **Table S1**).

#### Network anatomy and localization of circuits

##### Functional connectivity

At the node level, in the total sample, the highest strength nodes were located in the prefrontal cortex including the dorsolateral prefrontal cortex and cerebellum, showing distributed patterns of connections with sensorimotor, parietal, and other cerebellar nodes (**Figure 2**). The highest predictive nodes were within sensorimotor regions for males (**Figure 3**, top panel) and temporoparietal regions for females (**Figure 3**, bottom panel). Significant edges are visualized in **Figures 2-3**.

At the network level, reduced sensorimotor-cerebellum functional connectivity emerged as a consistent predictor of higher concurrent disruptive behavior severity in the total sample (**Figure 2**) as well as in male and female subgroups (**Figure 3**, top and bottom panels, respectively). Positive within-network sensorimotor connectivity also emerged as a shared predictor across males and females. In addition, positive connections between the medial frontal, salience, and sensorimotor networks and connections within the sensorimotor network were observed in the total sample (**Figure 2**) and male subgroup (**Figure 3**, top panel). Beyond these shared predictors, reduced sensorimotor connectivity with visual networks emerged as an important predictor of higher disruptive behavior severity in females (**Figure 3**, bottom panel).

##### Structural connectivity

In the total sample, the highest strength nodes were in the prefrontal cortex including the ventrolateral prefrontal cortex, temporoparietal, occipital, and cerebellar regions (**Figure 2**). In males, the highest strength nodes were focused in the right prefrontal and parietal cortex (**Figure 3**, top panel). In females, the hubs were sparser and distributed across prefrontal, cerebellar, occipital, subcortical, and limbic nodes showing the highest combined network strength (**Figure 3**, bottom panel). Significant edges are visualized in **Figures 2-3**.

At the network level, increased cerebellum-visual structural connectivity emerged as a prominent predictor of higher disruptive behavior severity in the total sample, alongside frontoparietal-visual association connectivity, with reduced salience-medial frontal connectivity contributing to the negative network (**Figure 2**). In males, increased frontoparietal-visual association connectivity was the greatest positive network level predictor, while reduced within network sensorimotor connectivity was the greatest negative network level predictor (**Figure 3**, top panel). In females, network level findings are not reported given the non-significance of the positive and negative networks (described above) (**Figure 3**, bottom panel).

### Prospective Models

#### Prediction results

##### Functional connectivity

Baseline functional connectivity predicted two-year change in disruptive behavior severity in the total sample (*ρ* = 0.038, RMSE = 4.425, *p_FDR_* = 0.027), with positive and negative edges contributing comparably (**Table S1**, **Figure 1**). In sex-stratified analyses, the combined and positive edge functional models were significant in males (combined: *ρ* =0.05, RMSE = 4.687, *p_FDR_* =0.027; positive edges: *ρ* = 0.05, RMSE = 4.683, *p_FDR_* = 0.023), whereas only the positive edge model was significant in females (positive edges: *ρ* = 0.029, RMSE = 4.154, *p_FDR_* = 0.045) (**Table S1**, **Figure 2**).

##### Structural connectivity

Baseline structural connectivity predicted two-year change in disruptive behavior severity in the total sample, which was driven by negative edges (ρ = 0.026, RMSE = 4.574, *pFDR* = 0.026) (**Table S1**, **Figure 1**). Combined and positive edge models were non-significant. Sex-stratified analyses revealed significant combined and negative edge models in males only (negative edges: ρ = 0.04, RMSE = 4.855, *pFDR* = 0.008). No prospective structural models reached significance in females (**Table S1**, **Figure 2**).

### Supplemental Follow-Up

As a sensitivity analysis, all CPM models were repeated with additional adjustment for CBCL Attention and Internalizing Problems subscale scores. Results were highly similar to the primary analyses across functional and structural connectivity models, with negligible changes in predictive performance. Importantly, the pattern of significant and non-significant findings remained unchanged following permutation testing and FDR correction (see **Supplemental Results**).

#### Network anatomy and localization of circuits

##### Functional connectivity

Prospective functional models included high-degree nodes in frontotemporal, medial frontal, supplementary motor, and parietal regions, with the dorsolateral prefrontal cortex prominent in males and the medial prefrontal cortex, supplementary motor area, and parietal cortex/angular gyrus prominent in females. The significant edges for within-and between-network interpretations are visualized in **Figure 2** (total sample) and **Figure 3** (top panel: males; bottom panel: females).

At the network level, increased baseline connectivity within the medial frontal network and between the medial frontal and sensorimotor networks predicted worsening disruptive behavior in the total sample (**Figure 2**) and males (**Figure 3**, top panel). In females, the strongest contributions were driven by increased connectivity between the frontoparietal and sensorimotor networks, and between the subcortical and salience networks (**Figure 3**, bottom panel).

##### Structural connectivity

In the total sample, the nodes with the highest degree negative connections were in occipital and subcortical (caudate, brainstem) regions. In males, the negative and combined networks shared a similar profile, with the highest strength nodes located in lateral prefrontal cortex including the inferior frontal gyrus, subcortical, and occipital regions. Significant edges are visualized in **Figures 2-3**.

At the network level, reduced baseline connectivity within and between frontoparietal, medial frontal, sensorimotor, subcortical, cerebellum, and visual networks emerged as prominent predictors of worsening disruptive behavior in the total sample (**Figure 2**). Reduced baseline sensorimotor-visual and cerebellum-salience connectivity also predicted increasing symptom severity across the two-year follow-up period in the total sample (**Figure 2**). In males, reduced baseline connectivity between frontoparietal-medial frontal, frontoparietal-visual, frontoparietal sensorimotor, medial frontal-subcortical, and cerebellum-visual connectivity were highly predictive of increasing disruptive behaviors, as well as reduced cerebellum-salience and subcortical-visual association connectivity (**Figure 3** top panel).

### Subnetwork level comparisons

Supplemental follow-up comparisons of significant CPM feature masks revealed significant model-specific contributions, particularly for functional vs structural connectivity and concurrent vs prospective functional models, and males vs females (all *p_FDR_* <0.032) (**Figure 4**; **Supplemental Results Tables S2-S3** and **Figures S1-S2**). No FDR-corrected comparisons were observed for shared connectivity patterns across modalities, time points, or sex-stratified subsamples (all *p_FDR_* >0.7), indicating that predictive models demonstrated partially distinct connectome configurations.

**Figure 4.**
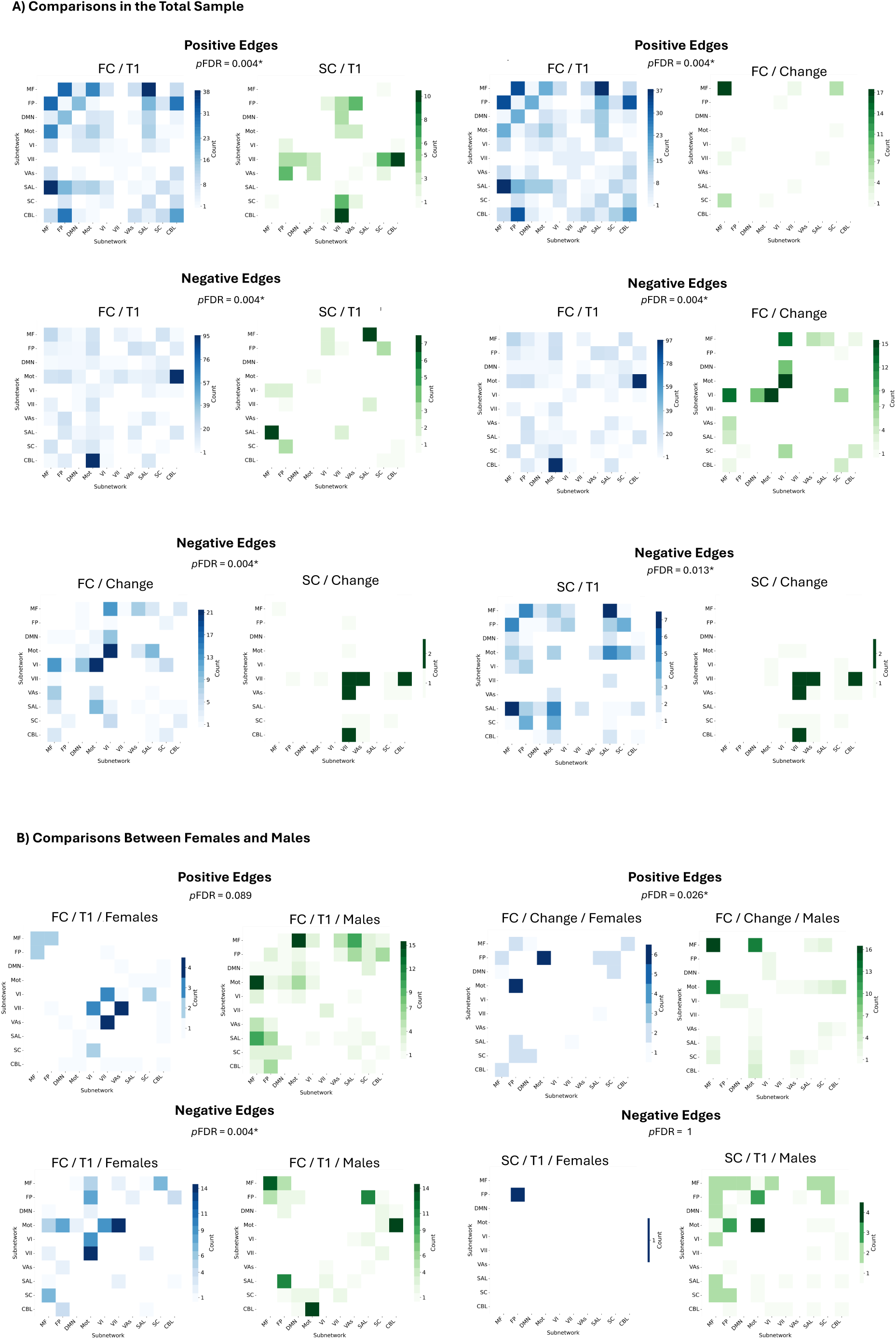
Supplemental follow-up of subnetwork-level comparison of CPM feature masks. A) Unique network pair comparisons of the total sample. B) Unique network pair comparisons for the female vs male samples. Network pairs that are unique in each comparison for the total sample, male subgroup, and female subgroup. The models being compared include functional connectivity (FC) vs structural connectivity (SC), cross-sectional or Time 1 (T1) vs prospective change (Change), and males vs females. Scores are reported separately for positive and negative networks. Benjamini-Hochberg false discovery rate correction (FDR) was applied to values. Network pairs are reported separately for positive and negative networks. Only comparisons reaching significance in the main CPM analyses are shown. The *p*FDR values correspond to Supplemental Results Table S2.

## Discussion

This study examined functional and structural connectome markers of concurrent disruptive behavior severity and two-year symptom change in a large, sex-balanced sample of youth. Three main findings emerged. First, both functional and structural connectivity predicted concurrent severity across distributed networks, whereas prospective associations were more focal. Functional connectivity predicted symptom change in the total sample, while prospective structural prediction was confined primarily to negative edges. Second, predictive connections involved distributed circuits including frontoparietal, medial frontal, sensorimotor, salience, default mode, cerebellar, subcortical, and sensorimotor networks. Notably, frontoparietal circuitry contributed across several models through lateral prefrontal and parietal nodes, including connections with visual association and sensorimotor networks. Third, predictive connectome markers showed partially distinct patterns across modality, time point, and sex-stratified subsamples, suggesting that functional and structural connectomes provide complementary and distinct information about disruptive behavior. Sex-stratified exploratory models showed partially distinct predictive networks across male and female subgroups (see **Supplemental Discussion**). Additionally, concurrent models showed broader predictive patterns and higher performance than prospective models, with limited overlap suggesting relatively distinct connectome markers of current symptom severity and subsequent symptom change (see **Supplemental Discussion**).

### Multimodal Predictive Performance and Connectivity Patterns

Both functional and structural connectomes predicted disruptive behavior severity. Functional connectivity tended to yield higher predictive estimates than structural connectivity, which is consistent with recent studies comparing functional and structural connectivity predicting cognitive [86] and behavioral outcomes [87], and general psychopathology [88]. However, a recent study found that functional connectivity was associated with general psychopathology, while structural connectivity, measured using fractional anisotropy, was associated with disruptive behavior severity [54]. Thus, our findings could suggest differences in the information gleaned from functional and structural connectomes, which supports complementary predictive contributions of each modality to identifying robust biomarkers specific to concurrent disruptive behavior and symptom course.

Regarding functional connectivity, predictive edges were distributed across canonical large-scale networks, with relatively balanced contributions from positive and negative networks. Concurrent models included high degree nodes in frontoparietal regions, particularly the dorsolateral prefrontal cortex, and cerebellum, whereas prospective models showed a more focal pattern of high degree nodes spanning frontoparietal, frontotemporal, medial frontal, sensorimotor, salience, and subcortical/limbic regions. At the network level, reduced sensorimotor-cerebellar connectivity was also a predictor of greater concurrent disruptive behavior. Additionally, increased baseline connectivity within the medial frontal network and between medial frontal and sensorimotor networks contributed to prospective prediction of increasing symptoms. Across concurrent and prospective models, frontoparietal and medial frontal networks emerged as highly predictive features, particularly for positive edges. The frontoparietal and medial frontal networks are associated with emotion regulation and cognitive control broadly [89-93], and implicated in disruptive behavior problems in youth [94,95]. Work from our group found consistent associations between emotion regulation difficulties and disruptive behavior severity [14], and a broader network dysfunction pattern linked to maladaptive aggression with frontoparietal network and related nodes, including the dorsolateral prefrontal cortex, emerging as highly predictive features [19]. Recent lesion mapping studies further support the role of frontoparietal nodes in maladaptive aggression and emotion regulation impairments, including the dorsolateral and ventrolateral prefrontal cortices [95-97]. Together, these findings suggest that frontoparietal circuitry may contribute to explaining heterogeneity in disruptive behavior through its interactions with medial frontal, sensorimotor, salience, limbic, and social perception networks.

Regarding structural connectivity, predictive edges were distributed across medial and lateral prefrontal, posterior parietal, temporoparietal, visual, subcortical, and cerebellar nodes. Concurrent structural models showed high degree nodes in the ventrolateral prefrontal cortex and frontoparietal regions. At the network level, increased frontoparietal-visual and cerebellar-visual connectivity contributed to greater concurrent severity, with frontoparietal connections emerging as the largest positive network level contributor in males. Prospective structural models revealed a more focal pattern spanning lateral prefrontal, occipital, subcortical, visual, cerebellar, and salience regions, and were driven primarily by negative edges. In the total sample, reduced baseline structural connectivity predicted greater subsequent symptom increases. In males, lateral prefrontal nodes, including the inferior frontal gyrus, contributed alongside occipital and subcortical regions, while reduced cerebellar-visual connectivity was also a contributing network level predictor. Prior work indicates disruptive behavior severity in youth is linked to altered microstructural integrity and white matter organization [54], particularly within association tracts, such as the superior longitudinal fasciculus and uncinate fasciculus, connecting frontoparietal and frontolimbic nodes involved in emotion regulation and broader cognitive control [35-41]. Our findings extend this prior work by identifying structural connectome patterns associated with concurrent severity and prospective symptom change. While the current findings indicate associations between the structural connectome and concurrent as well as worsening symptoms, given that quantitative anisotropy and other related microstructural metrics can be influenced by multiple aspects of tract and white matter organization, conclusions cannot be drawn related to network maturation. Future longitudinal studies in clinically affected youth will be important to understand connectome changes across development.

### Limitations

This study used baseline connectomes to predict symptom change. Future work will be important to simultaneously model both connectome and symptom changes over time. While the ABCD Study provides a rich population-level longitudinal dataset, neuroimaging is completed biannually, which could miss opportunities for capturing more rapid connectome and symptom changes during the transition from late childhood to early adolescence, a sensitive period of development. Future research could test whether persisting and desisting courses of disruptive behavior are associated with distinct circuit markers of multimodal connectivity. Predictive effect sizes were also small and modest (*ρ* = 0.026-0.087), with larger estimates for concurrent than prospective prediction. Importantly, these effect sizes are consistent with prior connectome studies in the ABCD Study and related transdiagnostic symptom domains [19,98], as well as brain-wide association studies in samples of similar size [86]. Predictive modeling approaches applied to large-scale datasets are also expected to produce smaller and more conservative estimates [98,99]. Additionally, smaller effect sizes may partly reflect the community-based ABCD Study sample, in which average disruptive behavior severity was relatively low and predominately within normative symptom ranges [100]. Relatively few youth in the clinically significant range may limit the ability of predictive models to capture the full heterogeneity of disruptive behavior. Future longitudinal neuroimaging studies in clinically affected youth will be important for evaluating connectome markers across a broader range of symptom severity. The current study focused on resting-state and structural connectivity, which do not directly probe the hypothesized emotion regulation circuitry. Future studies using explicit emotion regulation tasks, such as cognitive reappraisal, will be necessary to determine whether the identified frontoparietal and medial frontal circuits are directly implicated in disruptive behavior during explicit regulation and over time. Nonetheless, resting-state and structural MRI circumvent the cognitive demands of task fMRI and may improve imaging accessibility for youth across a range of symptom severity [70].

## Conclusion

The current findings may implicate multimodal connectomic markers in persistent disruptive behavior in youth and could inform future treatments, particularly during adolescence, which is a sensitive period of development marked by substantial neural reorganization and for risk of psychopathology. These findings suggest that concurrent disruptive behavior symptoms and later symptom worsening are linked to functional and structural connectivity markers in youth.

## Data Availability Statement

Data from the ABCD Study are shared on the NIH Brain Development Cohorts (NBDC) Data Hub https://www.nbdc-datahub.org. To promote transparency, all code used for analyses is available on GitHub (https://github.com/EmotionNeuroscienceLab).

## Acknowledgements

We would like to thank Ms. Olivia Ciocca for assisting with the diffusion MRI literature review. We also thank Dr. Matthew Rosenblatt for assistance with developing initial code. Claude (Anthropic) was used to assist in developing the R code for formatting the histograms in Figure 1 and in refining the R code for summarizing participant characteristics in Table 1. All AI-assisted code was reviewed by the authors, who take full responsibility for the figures and tables. The AI tool was used only for code generation assistance for Figure 1 and Table 1, and did not contribute to the study design, data interpretation, or scientific conclusions. A portion of the data used in the preparation of this article were obtained from the Adolescent Brain Cognitive Development (ABCD) Study (https://abcdstudy.org), held in the former NIMH Data Archive (NDA) and currently held in the NIH Brain Development Cohorts (NBDC) Data Hub. This is a multisite, longitudinal study designed to recruit more than 10,000 children ages 9-10 and follow them over 10 years into early adulthood. The ABCD Study® is supported by the National Institutes of Health and additional federal partners under award numbers U01DA041022, U01DA041028, U01DA041048, U01DA041089, U01DA041106, U01DA041117, U01DA041120, U01DA041134, U01DA041148, U01DA041156, U01DA041174, U24DA041123, U24DA041147, U01DA041093, and U01DA041025. A full list of supporters is available at https://abcdstudy.org/federal-partners.html. A listing of participating sites and a complete listing of the study investigators can be found at https://abcdstudy.org/consortium_members/. ABCD consortium investigators designed and implemented the study and/or provided data but did not necessarily participate in analysis or writing of this report. This manuscript reflects the views of the authors and may not reflect the opinions or views of the NIH or ABCD consortium investigators. The ABCD data repository grows and changes over time. The ABCD data used in this report came from NDA 5.1 release (DOI: https://doi.org/10.82525/wjkw-bp63). We are grateful to the study participants for their time and participation.

## Author Contributions

Conception: K.I., D.G.S., D.S.; Design of the work: K.I., D.G.S., E.C., D.S.; Acquisition, analysis, interpretation of data: K.I., D.G.S., E.C., C.L., Q.L., R.R., Y.Z., D.S.,; Manuscript writing and revising: K.I., D.G.S., E.C., K.M., C.L., Q.L., R.R., Y.Z., D.S.

