## Supplement for "Multimodal Connectome Markers of Disruptive Behavior Problems in Youth: Distinct Patterns of Concurrent Severity and Prospective Symptom Change"

**Supplemental Background:**

**Sex Differences in Network Markers of Disruptive Behavior**

Despite identified sex differences in network development related to connectivity strengthening [1-5] and cognitive control network maturation, particularly frontoparietal circuitry supporting emotion regulation [1-11], the majority of neuroimaging studies of childhood disruptive behavior have included predominantly male samples or lacked sufficient power to characterize connectivity patterns separately in males and females [12,13]. Prior work has reported mixed findings of sex differences in neuroanatomical markers of disruptive behavior problems in youth [14-16] and functional connectivity [17], while other studies have reported sex-by-diagnosis interactions in frontoparietal node neuroanatomy [18-21], white matter microstructure [22,23], and functional connectivity [24-28]. Work from our group in a sample of clinically affected youth with disruptive behavior problems reported sex differences in functional connectivity and neuroanatomy in nodes spanning frontolimbic and frontoparietal circuitry [19,26]. Another study reported sex differences in functional connectivity patterns linked to disruptive behavior in males and females, whereby later symptom changes was related to hyperconnectivity between the frontoparietal and default mode networks in males, but hypoconnectivity between default mode and limbic networks in girls [28]. Thus, findings are mixed and contradictory regarding sex differences in network connectivity in disruptive behavior in youth. Given the inconsistency of the existing evidence, the current study evaluates sex-stratified analyses as exploratory.

**Supplemental Methods:**

**Sample Selection**

The current study used data from the ABCD Study® (Data Release 5.1), which is a large-scale longitudinal study following 11,878 youth into early adulthood across 21 sites in the United States [29-32]. Data for the present study were from participants with available measures at baseline (9-10 years) and two-year follow-up (11-12 years). This subset consisted of participants with high-quality resting-state fMRI and diffusion weighted imaging (DWI) data. Of 11,878 enrolled participants, 10,988 had available resting-state fMRI and/or diffusion-weighted imaging data. We implemented selection criteria consistent with our prior work [33,34] and other ABCD studies, and included participants with low motion (i.e., mean framewise displacement of ≤0.25) and exclusion of connectomes with invalid regional data (i.e., non-zero edges for nodes). For participants with more than 1 run of resting-state, we selected the run with lowest motion for consistency across models. Participants were also required to have complete behavioral data at baseline and two-year follow-up. Our final sample included 7,556 participants with functional connectomes (3,726 females) and 8,978 participants with structural connectomes (4,343 females). See **Table 1** for participant characteristics.

**Behavioral Measures**

**Child Behavior Checklist (CBCL).** Externalizing behaviors were indexed using the Child Behavior Checklist (CBCL) [35], a continuous, transdiagnostic measure of child behavioral symptoms, to facilitate comparison with prior ABCD Imaging studies [34,36-39]. The CBCL is a parent-reported scale that indexed symptom severity across attention, externalizing and internalizing domains in the past six months. The CBCL Externalizing Problem subscale consists of the 12-item Rule-Breaking Behavior and 20-item Aggressive Behavior subscales, each rated on a 3-point Likert scale, and captures disruptive behaviors including verbal and physical aggression, conduct problems, anger/irritability, and noncompliance. The CBCL Externalizing Problem subscale was used as the primary measure of disruptive behavior problems. The CBCL Internalizing Problems scale includes items related to somatic complaints, social withdrawal, and anxiety/depressed symptoms. The CBCL Attention Problems scale includes symptoms related to inattention, hyperactivity, and impulsivity.

**Cognition.** Cognitive performance was evaluated using NIH Toolbox Cognition Battery [40]. Seven tasks, including the Picture Vocabulary (language), Flanker Test (attention), List Sort Working Memory Task (working memory), Dimensional Change Card Sort Task (executive function), Pattern Comparison Processing Speed Task (processing speed), Picture Sequence Memory Task (episodic memory), and the Oral Reading Test (language) were administered for 6 sub-domains (Weintraub, 2013). The age-corrected total composite score was used in analyses to regress out the influence of overall cognitive ability [40].

**ABCD Study: Imaging data acquisition and scan session descriptions**

The design and imaging protocol of the larger ABCD Study has been described in prior work [29]. Details of ABCD Study recruitment, neurocognitive batteries, and imaging protocols are also described elsewhere [29-32]. For more detail regarding the ABCD Study scan sequences and/or imaging acquisition parameters for structural (T1-weighted, T2-weighted), diffusion MRI, and fMRI, see Casey et al. [29] and Hagler et al. [30]. We also provide a summary here relevant to resting-state fMRI and diffusion-weighted imaging (DWI) modalities that were used in the current study. We used ABCD scan data including the structural T1-weighted series [30], diffusion MRI series using a multiband EPI [30,41,42] that includes 96 diffusion directions (6 directions at b=500 s/mm$^2$, 15 at b=1000 s/mm$^2$, 15 at b=2000 s/mm$^2$, and 60 at b=3000 s/mm$^2$), and resting-state series that includes at least 8 minutes of viewing a blank screen with a cross-hair with eyes open. For more detail regarding the ABCD Study scan sequences and acquisition parameters, see Casey et al. [29] and Hagler et al. [30].

**Preprocessing and Analysis**

**Resting-state fMRI preprocessing**. Functional connectomes were generated following an established pipeline (Barron et al., 2021). Images were motion-corrected in SPM12 and preprocessed in BioImage Suite. Nuisance covariates were regressed out, including linear and quadratic temporal drifts, mean signals from the cerebral spinal fluid (CSF), white matter, and gray matter, as well as a 24-parameter motion model (comprising six rigid-body motion parameters, their temporal derivatives, and all squared terms). Global signal regression was performed, given that it has been shown to improve brain-behavior associations and generalization of predictive models. Temporal filtering was applied using a Gaussian filter with an approximate cutoff of 0.12 Hz. The Shen 268-node atlas was used for brain parcellation [43]. For each subject, a 268×268 functional connectivity matrix was computed by the pairwise Pearson correlation between mean regional BOLD time series, followed by Fisher z-transformation to improve normality. Head motion parameters in resting-state fMRI were estimated through mean framewise displacement [44], a commonly used metric of head motion, which was used in connectome analyses as a continuous variable. Framewise displacement represents the sum of movement across the six rigid body motion parameters and across a scan series and was included in subsequent connectome analyses to account for residual motion effects.

**Diffusion-Weighted Imaging (DWI) Preprocessing.** Diffusion-weighted images were preprocessed using the SHARD pipeline and reconstructed using generalized q-sampling imaging. Whole-brain fiber tracking was then performed in DSI Studio, generating 1,000,000 streamlines per subject [45]. Structural connectivity matrices were derived by registering each participant's brain to a standard space and computing the pairwise connectivity strength between regions. Connectivity was quantified using the mean quantitative anisotropy value of streamlines connecting each ROI pair [46]. A threshold of 0.001 was applied to exclude spurious connections. The resulting structural connectome was a 268×268 weighted adjacency matrix representing the anatomical connectivity between all regions defined by the same Shen atlas. Masked diffusion-weighted imaging correlation estimates were used as a proxy for motion and image-quality covariate, consistent with ABCD diffusion MRI quality-control procedures [30].

The masked diffusion-weighted imaging correlation reflects the similarity of diffusion-weighted volumes after brain masking and provides an index of motion and image-quality variation in diffusion MRI data.

**Generating Multimodal Connectomes**

**Functional Connectomes**. Resting-state fMRI data were preprocessed using an established pipeline implemented in SPM12 and BioImage Suite [47-49], consistent with prior connectome-based predictive modeling studies [34]. Preprocessing included motion correction, registration to each participant's anatomical image, nuisance regression, global signal regression, and temporal filtering (using a Gaussian filter with an approximate cutoff of 0.12 Hz). Global signal regression was performed, given that it has been shown to improve brain-behavior associations and generalization of predictive models. Nuisance regressors included linear and quadratic temporal trends, mean cerebrospinal fluid signal, mean white matter signal, mean gray matter signal, and a 24-parameter motion model comprising six rigid-body motion parameters, their temporal derivatives, and squared terms. Mean BOLD time series were extracted from each Shen atlas region [43]. Functional connectivity was calculated as the pairwise Pearson correlation between mean regional BOLD time series, and correlation coefficients were Fisher z-transformed prior to analysis. This procedure yielded one 268 × 268 functional connectivity matrix for each participant. For functional connectivity analyses, motion was quantified using mean framewise displacement [44], consistent with established resting-state fMRI motion-control procedures [50].

**Structural Connectome.** Diffusion-weighted images were preprocessed using the SHARD pipeline and reconstructed using generalized q-sampling imaging in DSI Studio [45,46]. Whole-brain deterministic fiber tracking was performed in DSI Studio, generating 1,000,000 streamlines per participant. The Shen atlas was registered to each participant's diffusion/anatomical space and used to define regions of interest for structural connectome construction. Structural connectivity between each pair of regions was quantified as the mean quantitative anisotropy value of streamlines connecting the two regions [45,46]. Quantitative anisotropy provides an estimate of the peak density of water diffusion along the main axes of white matter fibers as a proxy for microstructural integrity and connectivity. Higher quantitative anisotropy reflects higher structural connectivity and white matter microstructural integrity. Connections with weights below 0.001 were removed to reduce weak or potentially spurious connections. The resulting structural connectome was a 268 × 268 weighted adjacency matrix representing anatomical connectivity among the same Shen atlas regions used in the functional connectivity analysis. In the current study, we use quantitative anisotropy, which is derived from q-space imaging and provides sensitivity to microstructural variation, because this metric is less susceptible to partial volume effects and free water contamination, and has shown greater reproducibility compared with conventional diffusion-weighted imaging measures [45,46] and has been used in prior studies [51-53], including connectomics work from our group [54-56]. For structural connectivity analyses, masked neighboring diffusion-weighted imaging correlation (NDC) was used as a motion and image-quality covariate [57], consistent with ABCD diffusion MRI quality-control procedures [30]. This measure reflects the similarity of diffusion-weighted volumes after brain masking and provides an index of motion-related artifacts or poor data quality variation in diffusion MRI data [57].

**Connectome-based Predictive Modeling**

Connectome-based predictive modeling (CPM) was used to test whether whole-brain functional and structural connectivity patterns predicted disruptive behavior [47-49,58]. CPM was implemented separately for functional and structural connectivity matrices. Models were constructed to predict baseline disruptive behavior and longitudinal symptom change over two years. CPM was implemented using 10-fold cross-validation repeated 100 times to estimate out-of-sample predictive performance. Within each training fold, partial correlations were calculated between each connectivity edge and the behavioral outcome, adjusting for model-specific covariates. Edges positively associated with the outcome at $p \leq 0.01$ were retained in a positive feature mask, and edges negatively associated with the outcome at $p \leq 0.01$ were retained in a negative feature mask. Feature selection was performed only within the training folds to prevent information leakage [49,58,59].

For each participant, positive and negative network strength scores were calculated by summing connectivity weights across edges in the corresponding feature masks. For the combined model, network strength was calculated as positive network strength minus negative network strength and entered as a single predictor in a robust linear regression model. Positive and negative edge models were fitted separately using robust linear regression and the corresponding network strength scores. Training-derived feature masks and fitted model coefficients were applied to held-out participants to generate predicted behavioral scores. Thus, three model types were evaluated separately: a combined model incorporating both positive and negative edges, a positive edge model, and a negative edge model. Predictive performance was quantified as the Spearman correlation between observed and predicted behavioral scores across held-out participants. Spearman correlations between observed and predicted behavioral scores were used as the primary performance estimates. For each model type, the repetition with the median-ranked prediction performance was selected, and the corresponding Spearman correlation was retained for reporting.

Statistical significance was assessed using permutation testing with 1,000 permutations. For each permutation, behavioral labels were randomly shuffled and the full CPM procedure was repeated to generate a null distribution of prediction accuracy. Empirical permutation *p* values were calculated as the proportion of permuted models with prediction accuracy greater than or equal to the observed prediction accuracy. Permutation-derived *p* values were subsequently adjusted for multiple comparisons using the Benjamini-Hochberg false discovery rate procedure.

**Identification of Disruptive Behavior Connectomes**

The CBCL Externalizing Behavior Problems subscale score [35] was used as the primary dimensional measure of disruptive behavior severity in predictive models. For cross-sectional models, baseline CBCL Externalizing Problems raw scores were used as the dependent variable. For prospective models, CBCL Externalizing Problems symptom change was calculated as the two-year follow-up score minus the baseline score. Higher values therefore indicated greater symptom increases or less symptom reduction over time. Prospective models additionally adjusted for baseline disruptive behavior severity. Baseline functional and structural connectivity were used as independent variables in modality-specific CPM analyses. Models were conducted in the full sample and then in sex-stratified analyses conducted separately in males and females.

Covariate-adjusted models included age, cognitive performance, and in-scanner head motion. Cognitive performance was measured using the NIH Toolbox Cognition Battery total composite score [40]. Sex was included as an additional covariate in full-sample models but was not included in sex-stratified models. Functional connectivity models included mean framewise displacement as the motion covariate [44]. Structural connectivity models included masked diffusion-weighted imaging correlation as the diffusion MRI motion and image-quality covariate. Prospective models additionally controlled for baseline disruptive behavior severity to test whether baseline connectivity predicted subsequent symptom change beyond initial symptom burden.

**Subnetwork-Level Comparison of CPM Feature Masks**

To assess the extent to which CPM-derived predictive networks showed shared versus model-specific topographic organization, feature masks were compared at the subnetwork level. CPM-selected edge masks were first mapped from the Shen 268-node parcellation to ten canonical subnetworks using the predefined Shen subnetwork assignments. For each mask, selected edges were aggregated into a 10×10 subnetwork count matrix, with each cell representing the number of selected edges within or between a given pair of subnetworks.

Within-subnetwork connections were retained in these summaries. For each pairwise comparison, we quantified three subnetwork-level metrics: the number of subnetwork pairs selected in both masks, the number of subnetwork pairs uniquely selected in the first mask, and the number of subnetwork pairs uniquely selected in the second mask. A subnetwork pair was considered present in a given mask if its corresponding subnetwork-level count was greater than zero. Statistical significance was assessed using non-parametric permutation testing with 1,000 iterations. For each comparison, the subnetwork-level counts from the two masks were pooled, randomly reassigned into two permuted masks of equal size, and used to generate empirical null distributions for shared and unique subnetwork-pair counts. Empirical p-values were calculated as the proportion of permuted values greater than or equal to the observed statistic. Analyses were restricted to CPM models that demonstrated significant predictive performance. Thus, pairwise comparisons were performed separately by imaging modality, time point, sample subgroup, and edge polarity. The follow-up contrasts included functional and structural connectivity models (edges) that emerged as significant in predicting baseline disruptive behavior and prospective change.

**Terminology Referring to Disruptive Behavior Problems**

The term *disruptive behavior problem* is used as a transdiagnostic construct encompassing behaviors spanning harm to self or others, noncompliance, defiance, rule-breaking behaviors, and irritability/anger. While disruptive behavior problems can also be conceptualized as symptom subtypes, this study focuses on the disruptive behavior spectrum among children and adolescents whose families were seeking outpatient treatment for their child's disruptive behavior problems [60].

Related constructs of disruptive behavior problems include externalizing problems and maladaptive aggression. Disruptive behavior problems (or externalizing problems) refer to symptoms including irritability/anger, aggression, noncompliance, and rule-breaking behavior. Maladaptive aggression is one component of disruptive behavior and includes impulsive, affective, hostile, and reactive forms of aggression. Maladaptive aggression can also reflect an emotional response to frustration or provocation and may manifest as anger outbursts, yelling, screaming, pushing, hitting, or throwing objects [61,62].

We used the parent-reported Child Behavior Checklist (CBCL) Externalizing Behavior Problems subscale, which is a broad band measure of disruptive behavior problems, as a continuous, transdiagnostic measure of disruptive behavior severity. The CBCL Externalizing Problems scale comprises the Aggressive Behavior (20 items) and Rule-Breaking Behavior (12 items) subscales, with items rated on a 3-point Likert scale. The Externalizing Problems scale assesses a broad range of disruptive behaviors, including physical and verbal aggression, conduct problems, irritability/anger, and noncompliance. The term disruptive behavior disorders is used to reflect the DSM-5 diagnostic classification including oppositional defiant and conduct disorders, which are characterized by patterns of anger/irritability, defiance, vindictiveness, and/or oppositional behavior [63,64].

**Supplemental Results:**

**Sensitivity analyses**

As a supplemental sensitivity analysis, all CPM models were repeated with additional adjustment for baseline CBCL Attention and Internalizing Problems subscale scores. Results showed a highly similar pattern of predictive performance and significance to the primary analyses, indicating that findings were not attributable to co-occurring attention or internalizing symptom severity. Across modalities, inclusion of attention and internalizing symptoms as covariates resulted in negligible changes in significance patterns including similar effect sizes and permutation-derived, FDR-corrected *p* values.

For functional connectivity, cross-sectional models remained significant in the total sample (all $\rho$ = 0.080-0.087, all $p_{FDR} < 0.004$), males (all $\rho$ = 0.065-0.085, all $p_{FDR} < 0.009$), and females (all $\rho$ = 0.032-0.039, all $p_{FDR} \leq 0.03$). Prospective functional connectivity models also showed a highly similar pattern to the primary analyses, with significant predictions remaining in the total sample ($\rho$ = 0.034-0.036, $p_{FDR} \leq 0.04$), males for the combined and positive models (all $\rho$ = 0.053-0.055, $p_{FDR} < 0.03$), and females for the positive edge model ($\rho = 0.029$, $p_{FDR} < 0.05$).

Structural connectivity findings also remain highly similar to primary models. Cross-sectional models remained significant in the total sample (all $\rho$ = 0.023-0.071, all $p_{FDR} < 0.004$), males (all $\rho$ = 0.02-0.05, all $p_{FDR} < 0.034$), and females for the combined model ($\rho = 0.05$, $p_{FDR} = 0.023$). Prospective structural connectivity results closely replicated the primary analyses, with significant negative edge models in the total sample ($\rho = 0.026$, $p_{FDR} = 0.027$) and significant combined and negative edge models in males (all $\rho$ = 0.045-0.045, all $p_{FDR} < 0.008$), whereas female models remained non-significant (all $p_{FDR} > 0.5$).

**Table S1.** Multimodal Predictive Model Results and Performance

| Imaging Modality | Feature Type | $\rho$[a] | *RMSE*[b] | $p_{(perm)}$[c] | $p_{FDR}$[d] |
|---|---|---|---|---|---|
| **Functional Connectivity (resting-state fMRI)** | | | | | |
| *Cross-Sectional Models* | | | | | |
| Total Sample | Combined | 0.087 | 5.660 | < 0.001 | **< 0.001** |
| | Positive | 0.082 | 5.664 | < 0.001 | **< 0.001** |
| | Negative | 0.080 | 5.662 | < 0.001 | **< 0.001** |
| Males | Combined | 0.082 | 6.158 | < 0.001 | **< 0.001** |
| | Positive | 0.085 | 6.150 | < 0.001 | **< 0.001** |
| | Negative | 0.065 | 6.170 | 0.003 | **0.009** |
| Females | Combined | 0.038 | 5.039 | 0.017 | **0.028** |
| | Positive | 0.040 | 5.042 | 0.014 | **0.028** |
| | Negative | 0.032 | 5.039 | 0.024 | **0.032** |
| *Prospective Models* | | | | | |
| Total Sample | Combined | 0.038 | 4.425 | 0.013 | **0.028** |
| | Positive | 0.029 | 4.426 | 0.025 | **0.032** |
| | Negative | 0.033 | 4.425 | 0.024 | **0.032** |
| Males | Combined | 0.053 | 4.688 | 0.016 | **0.028** |
| | Positive | 0.055 | 4.683 | 0.009 | **0.023** |
| | Negative | 0.038 | 4.691 | 0.085 | 0.096 |
| Females | Combined | 0.023 | 4.164 | 0.102 | 0.107 |
| | Positive | 0.029 | 4.155 | 0.038 | **0.046** |
| | Negative | 0.023 | 4.163 | 0.107 | 0.107 |
| **Structural Connectivity (quantitative anisotropy)** | | | | | |
| *Cross-Sectional Models* | | | | | |
| Total Sample | Combined | 0.071 | 5.801 | < 0.001 | **< 0.01** |
| | Positive | 0.059 | 5.815 | < 0.001 | **< 0.01** |
| | Negative | 0.023 | 5.802 | < 0.001 | **< 0.01** |
| Males | Combined | 0.052 | 6.310 | < 0.001 | **< 0.01** |
| | Positive | 0.044 | 6.325 | < 0.001 | **< 0.01** |
| | Negative | 0.020 | 6.333 | 0.021 | **0.034** |
| Females | Combined | 0.054 | 5.142 | 0.012 | **0.024** |
| | Positive | 0.052 | 5.141 | 0.078 | 0.096 |
| | Negative | -0.004 | 5.128 | 0.010 | **0.024** |
| *Prospective Models* | | | | | |
| Total Sample | Combined | 0.022 | 4.576 | 0.050 | 0.069 |
| | Positive | 0.012 | 4.571 | 0.045 | 0.067 |
| | Negative | 0.026 | 4.575 | 0.015 | **0.027** |

| | | | | | |
|---|---|---|---|---|---|
| Males | Combined | 0.045 | 4.865 | 0.012 | **0.024** |
| | Positive | 0.020 | 4.867 | 0.080 | 0.096 |
| | Negative | 0.045 | 4.856 | 0.003 | **0.009** |
| Females | Combined | -0.023 | 4.269 | 0.913 | 0.967 |
| | Positive | -0.013 | 4.241 | 0.494 | 0.555 |
| | Negative | -0.027 | 4.277 | 1.000 | 1.000 |

*Note*: Spearman correlation coefficients between actual versus predicted CBCL Externalizing Behavior Problems severity scores. Correlations were computed separately for the total, male-only and female-only samples, each based on the combined, positive-only and negative-only edges.

[a]Spearman Correlation Coefficient

[b]Root Mean Square Error

[c]permutation *p*-value

[d]permutation *p*-value adjusted for multiple comparisons using the Benjamini-Hochberg false discovery rate procedure

**Table S2.** Supplementary Follow-Up of Subnetwork Pairs with at Least One Unique Edge

| **Modality/ Timepoint A** | **Modality/ Timepoint B** | **Group** | **Positive or Negative Edges** | **Shared Edges Score** | ***p* Shared Edges** | **$p_{FDR}$ Shared Edges** | **Unique Edges Score for A** | **Unique Edges Score for B** | ***p* Unique** | **$p_{FDR}$ Unique** |
|---|---|---|---|---|---|---|---|---|---|---|
| FC/T1 | SC/T1 | Total Sample | negative | 48 | 1 | 1 | 375 | 20 | **0.001** | **0.004** |
| FC/T1 | SC/T1 | Total Sample | positive | 82 | 1 | 1 | 334 | 44 | **0.001** | **0.004** |
| FC/T1 | SC /T1 | Females | negative | 1 | 1 | 1 | 102 | 3 | **0.012** | **0.032** |
| FC/T1 | SC/T1 | Males | negative | 17 | 1 | 1 | 87 | 19 | **0.001** | **0.004** |
| FC/T1 | SC /T1 | Males | positive | 43 | 1 | 1 | 103 | 38 | **0.001** | **0.004** |
| FC/Change | SC/Change | Total Sample | negative | 8 | 1 | 1 | 96 | 13 | **0.001** | **0.004** |
| FC/T1 | FC/Change | Total Sample | negative | 46 | 1 | 1 | 377 | 58 | **0.001** | **0.004** |
| FC/T1 | FC/Change | Total Sample | positive | 35 | 1 | 1 | 381 | 28 | **0.001** | **0.004** |
| FC/T1 | FC/Change | Females | positive | 18 | 0.985 | 1 | 65 | 13 | **0.046** | 0.105 |
| FC/T1 | FC/Change | Males | positive | 35 | 1 | 1 | 111 | 31 | **0.001** | **0.004** |
| SC/T1 | SC/Change | Total Sample | negative | 7 | 1 | 1 | 61 | 14 | **0.004** | **0.013** |
| SC /T1 | SC/Change | Males | negative | 13 | 0.741 | 1 | 23 | 19 | 0.396 | 0.791 |
| FC/T1/Females | FC/T1/Males | Females/ Males | negative | 45 | 1 | 1 | 58 | 59 | **0.001** | **0.004** |
| FC/T1/Females | FC/T1/Males | Females/Males | positive | 59 | 0.979 | 1 | 24 | 87 | **0.036** | 0.089 |
| FC/Change/Females | FC/Change/ Males | Females/Males | positive | 13 | 0.997 | 1 | 18 | 53 | **0.009** | **0.026** |
| SC/T1/Females | SC/T1/Males | Females/Males | negative | 3 | 0.381 | 0.791 | 1 | 33 | 0.9 | 1 |

*Note*: Subnetwork-level count summary statistics for shared and unique edge contributions across paired CPM models. For each comparison, predictive edge masks from the two models were reduced from the ROI-by-ROI level into subnetwork-by-subnetwork count matrices. The two models being compared are represented as A and B, which includes functional connectivity (FC) vs structural connectivity (SC), cross-sectional or Time 1 (T1) vs prospective change score (Change), and males vs females. Shared edges score reflects the total magnitude (i.e., summed count) of edges common to both models. Unique edges score for A and B reflect the magnitude specific to each. Scores are reported separately for positive and negative networks. Benjamini-Hochberg false discovery rate correction (FDR) was applied across the full set of 32 tests (16 shared and 16 unique p-values), with adjusted values reported as FDR. Only comparisons reaching significance in the primary CPM analyses are shown.

**Table S3.** Supplementary Follow-Up of Subnetwork Pairs Mapped onto Macroscale Networks

| Modality/ Timepoint A | Modality/ Timepoint B | Group | Positive or Negative Edges | Shared Network Pairs | Unique Network Pairs of A | Unique Network Pairs of B |
|---|---|---|---|---|---|---|
| FC/T1 | SC/T1 | Total Sample | negative | MF-MF, MF-FP, MF-DMN, MF-Mot, MF-SC, FP-FP, FP-DMN, FP-Mot, FP-VI, FP-SAL, FP-SC, DMN-SAL, DMN-CBL, Mot-Mot, Mot-VI, Mot-VAs, Mot-SAL, Mot-SC, Mot-CBL, SAL-SAL, SAL-SC, SAL-CBL | MF-MF, MF-FP, MF-DMN, MF-Mot, MF-VII, MF-VAs, MF-SC, MF-CBL, FP-FP, FP-DMN, FP-Mot, FP-VAs, FP-SAL, FP-CBL, DMN-DMN, DMN-Mot, DMN-VI, DMN-VII, DMN-SAL, DMN-SC, Mot-VI, Mot-VII, Mot-VAs, Mot-SAL, Mot-SC, Mot-CBL, VI-VI, VI-SAL, VI-SC, VI-CBL, VAs-VAs, VAs-SAL, VAs-SC, SAL-CBL, SC-SC | MF-VI, MF-SAL, FP-VI, FP-VII, FP-SC, Mot-Mot, VII-SAL, SC-CBL, CBL-CBL |
| FCI/T1 | SC/T1 | Total Sample | positive | MF-MF, MF-FP, MF-DMN, MF-Mot, MF-VI, MF-VAs, MF-SC, MF-CBL, FP-FP, FP-DMN, FP-Mot, FP-VAs, FP-SAL, FP-SC, FP-CBL, DMN-Mot, DMN-VII, DMN-VAs, DMN-SAL, DMN-SC, DMN-CBL, Mot-Mot, Mot-SAL, Mot-CBL, VI-VII, VI-SAL, VI-SC, VII-VII, VII-VAs, VII-CBL, VAs-SC, VAs-CBL, SAL-SC, SAL-CBL, SC-SC, SC-CBL | MF-FP, MF-DMN, MF-Mot, MF-VI, MF-VAs, MF-SAL, MF-CBL, FP-FP, FP-DMN, FP-Mot, FP-SAL, FP-SC, FP-CBL, DMN-Mot, DMN-VI, DMN-VAs, DMN-SAL, DMN-CBL, Mot-Mot, Mot-VI, Mot-SAL, Mot-SC, Mot-CBL, VI-VI, VI-VII, VI-SAL, VI-SC, VII-VII, VII-VAs, VAs-SAL, VAs-CBL, SAL-SAL, SAL-SC, SAL-CBL, SC-CBL, CBL-CBL | MF-SC, FP-VI, FP-VII, FP-VAs, DMN-VII, Mot-VII, Mot-VAs, VI-CBL, VII-SAL, VII-SC, VII-CBL, VAs-VAs, VAs-SC |

| | | | | | | |
|---|---|---|---|---|---|---|
| FC/T1 | SC/T1 | Females | negative | MF-Mot | MF-MF, MF-DMN, MF-Mot, MF-VII, MF-SAL, MF-SC, FP-Mot, FP-VAs, FP-SAL, FP-CBL, DMN-Mot, DMN-SC, Mot-VI, Mot-VII, Mot-VAs, Mot-SAL, Mot-SC, Mot-CBL, VI-VI, VII-SAL, VAs-SAL, SAL-CBL | MF-FP, FP-FP, FP-SC |
| FC/T1 | SC/T1 | Males | negative | MF-MF, MF-FP, MF-DMN, MF-Mot, MF-SC, FP-DMN, FP-Mot, FP-VAs, FP-SC, Mot-SAL, Mot-SC, SAL-CBL | MF-MF, MF-FP, MF-Mot, MF-VII, MF-SC, FP-FP, FP-DMN, FP-SAL, DMN-SC, DMN-CBL, Mot-VI, Mot-VII, Mot-VAs, Mot-SAL, Mot-SC, Mot-CBL, VI-VI, VI-VAs, VI-SAL, VII-SC, VAs-SAL | MF-DMN, MF-VI, MF-SAL, FP-Mot, FP-SC, FP-CBL, Mot-Mot, VI-CBL, VII-SAL, VII-CBL, SAL-SAL, CBL-CBL |
| FC/T1 | SC/T1 | Males | positive | MF-DMN, MF-Mot, MF-VAs, MF-SAL, MF-SC, FP-FP, FP-DMN, FP-Mot, FP-VAs, FP-SAL, FP-SC, FP-CBL, DMN-VAs, DMN-SAL, DMN-SC, DMN-CBL, Mot-Mot, Mot-VI, Mot-VAs, Mot-SAL, Mot-CBL, VI-SAL, VAs-CBL, SAL-CBL, SC-CBL | MF-FP, MF-DMN, MF-Mot, MF-VI, MF-VAs, MF-SAL, MF-SC, MF-CBL, FP-DMN, FP-SAL, FP-SC, FP-CBL, DMN-DMN, DMN-Mot, DMN-VI, DMN-SAL, Mot-Mot, Mot-VI, Mot-SAL, Mot-SC, VII-VII, VAs-SAL, SAL-SAL, SAL-SC, SC-CBL | FP-VI, FP-VII, FP-VAs, DMN-CBL, Mot-VII, Mot-VAs, VI-VII, VI-VAs, VI-CBL, VII-VAs, VII-SAL, VII-SC, VII-CBL, VAs-VAs, VAs-SC, VAs-CBL, SC-SC |
| FC/Change | SC/Change | Total Sample | negative | MF-Mot, Mot-VI, VI-SC, VAs-VAs, SAL-CBL, SC-CBL | MF-DMN, MF-VI, MF-VII, MF-VAs, MF-SAL, MF-CBL, FP-DMN, FP-VI, FP-SC, DMN-Mot, DMN-VI, Mot-Mot, Mot-VI, Mot-VAs, Mot-SAL, VI-SAL, VI-SC, VAs-SAL, SAL-SC, SAL-CBL, SC-CBL | MF-MF, FP-VII, Mot-VII, VII-VII, VII-VAs, VII-SC, VII-CBL, VAs-SC, VAs-CBL, SC-SC |

| | | | | | | |
|---|---|---|---|---|---|---|
| FC/T1 | FC/Change | Total Sample | negative | MF-DMN, MF-Mot, MF-VII, MF-VAs, MF-CBL, FP-DMN, FP-VI, FP-SC, DMN-Mot, DMN-VI, Mot-Mot, Mot-VI, Mot-VAs, Mot-SAL, VI-SAL, VI-SC, VAs-VAs, VAs-SAL, SAL-SC, SAL-CBL | MF-MF, MF-FP, MF-DMN, MF-Mot, MF-VII, MF-SC, FP-FP, FP-DMN, FP-Mot, FP-VAs, FP-SAL, FP-CBL, DMN-DMN, DMN-Mot, DMN-VII, DMN-SAL, DMN-SC, DMN-CBL, Mot-Mot, Mot-VII, Mot-VAs, Mot-SAL, Mot-SC, Mot-CBL, VI-VI, VI-SAL, VI-CBL, VAs-SAL, VAs-SC, SAL-SAL, SAL-CBL, SC-SC | MF-VI, MF-VAs, MF-SAL, MF-CBL, FP-SC, DMN-VI, Mot-VI, VI-SC, SC-CBL |
| FC/T1 | FC/Change | Total Sample | positive | MF-MF, MF-DMN, MF-Mot, MF-SAL, MF-SC, MF-CBL, FP-DMN, FP-Mot, FP-SAL, DMN-Mot, DMN-VI, DMN-CBL, Mot-SAL, Mot-SC, Mot-CBL, VI-VII | MF-FP, MF-DMN, MF-Mot, MF-VI, MF-VAs, MF-SAL, MF-CBL, FP-FP, FP-DMN, FP-Mot, FP-VAs, FP-SAL, FP-SC, FP-CBL, DMN-Mot, DMN-VI, DMN-VII, DMN-VAs, DMN-SAL, DMN-SC, DMN-CBL, Mot-Mot, Mot-VI, Mot-SAL, Mot-CBL, VI-VI, VI-VII, VI-SAL, VI-SC, VII-VII, VII-VAs, VII-CBL, VAs-SAL, VAs-SC, VAs-CBL, SAL-SAL, SAL-SC, SAL-CBL, SC-SC, SC-CBL, CBL-CBL | MF-MF, MF-VII, MF-SC, FP-VI, Mot-SC, VII-SAL |
| FC/T1 | FC/Change | Females | positive | MF-MF, MF-FP, MF-DMN, MF-Mot, MF-SAL, MF-SC, MF-CBL, FP-DMN, FP-Mot, FP-SAL, FP-SC, DMN-CBL, Mot-CBL | MF-FP, MF-Mot, MF-SAL, FP-DMN, FP-VII, FP-CBL, DMN-VAs, DMN-SAL, Mot-Mot, Mot-VAs, Mot-SAL, Mot-SC, Mot-CBL, VI-VII, VI-VAs, VI-SC, VII-VAs, VII-CBL, | MF-DMN, FP-Mot, FP-SAL, FP-SC, DMN-Mot, DMN-SC, VII-SAL |

| | | | | | | |
|---|---|---|---|---|---|---|
| | | | | | VAs-CBL, SAL-SC, SAL-CBL, SC-CBL, CBL-CBL | |
| FC/T1 | FC/Change | Males | positive | MF-DMN, MF-Mot, MF-SAL, MF-SC, FP-DMN, FP-Mot, DMN-Mot, DMN-VI, DMN-CBL, Mot-VAs, Mot-SAL, Mot-SC, Mot-CBL, VAs-CBL, SAL-SC | MF-FP, MF-DMN, MF-Mot, MF-VI, MF-VAs, MF-SAL, MF-CBL, FP-FP, FP-DMN, FP-Mot, FP-VAs, FP-SAL, FP-SC, FP-CBL, DMN-DMN, DMN-Mot, DMN-VAs, DMN-SAL, DMN-SC, Mot-Mot, Mot-VI, Mot-SAL, VI-SAL, VII-VII, VAs-SAL, SAL-SAL, SAL-SC, SAL-CBL, SC-CBL | MF-MF, FP-VI, DMN-VI, DMN-CBL, Mot-VII, Mot-SC, Mot-CBL, VAs-SC |
| SC/T1 | SC/Change | Total Sample | negative | MF-MF, MF-Mot, FP-VII, Mot-VI, SAL-CBL, SC-CBL | MF-MF, MF-FP, MF-DMN, MF-Mot, MF-VI, MF-SAL, MF-SC, FP-FP, FP-DMN, FP-Mot, FP-VI, FP-SAL, FP-SC, DMN-SAL, DMN-CBL, Mot-Mot, Mot-VAs, Mot-SAL, Mot-SC, Mot-CBL, VII-SAL, SAL-SAL, SAL-SC, SAL-CBL, CBL-CBL | Mot-VI, Mot-VII, VI-SC, VII-VII, VII-VAs, VII-SC, VII-CBL, VAs-VAs, VAs-SC, VAs-CBL, SC-SC |
| SC/T1 | SC/Change | Males | negative | MF-FP, MF-Mot, MF-SAL, MF-SC, FP-Mot, FP-SC, VII-SAL, VII-CBL, SAL-SAL, SAL-CBL | MF-MF, MF-FP, MF-DMN, MF-Mot, MF-VI, MF-SAL, FP-DMN, FP-Mot, FP-VAs, FP-SC, FP-CBL, Mot-Mot, Mot-SAL, Mot-SC, VI-CBL, CBL-CBL | FP-FP, FP-VII, VI-SC, VII-VAs, VII-SC, VII-CBL, VAs-VAs, VAs-SC, SAL-SC, SAL-CBL, SC-SC |

| | | | | | | |
|---|---|---|---|---|---|---|
| FC/T1/Females | FC/T1/Males | Females/ Males | negative | MF-MF, MF-DMN, MF-Mot, MF-VII, MF-SC, FP-Mot, FP-VAs, FP-SAL, DMN-SC, Mot-VI, Mot-VII, Mot-VAs, Mot-SAL, Mot-SC, Mot-CBL, VI-VI, VAs-SAL, SAL-CBL | MF-DMN, MF-Mot, MF-VII, MF-SAL, MF-SC, FP-Mot, FP-VAs, FP-CBL, DMN-Mot, Mot-VI, Mot-VII, VII-SAL, SAL-CBL | MF-MF, MF-FP, FP-FP, FP-DMN, FP-SAL, FP-SC, DMN-SC, DMN-CBL, Mot-SC, Mot-CBL, VI-VAs, VI-SAL, VII-SC, VAs-SAL |
| FC/T1/Females | FC/T1/Males | Females/ Males | positive | MF-FP, MF-DMN, MF-Mot, MF-SAL, MF-SC, MF-CBL, FP-DMN, FP-Mot, FP-SAL, FP-SC, FP-CBL, DMN-VAs, DMN-SAL, DMN-CBL, Mot-Mot, Mot-VAs, Mot-SAL, Mot-SC, Mot-CBL, VAs-CBL, SAL-SC, SAL-CBL, SC-CBL | MF-MF, MF-FP, FP-VII, DMN-VAs, DMN-CBL, Mot-SAL, Mot-SC, Mot-CBL, VI-VII, VI-VAs, VI-SC, VII-VAs, VII-CBL, VAs-CBL, SAL-CBL, CBL-CBL | MF-DMN, MF-Mot, MF-VI, MF-VAs, MF-SAL, MF-SC, MF-CBL, FP-FP, FP-DMN, FP-Mot, FP-VAs, FP-SAL, FP-SC, FP-CBL, DMN-DMN, DMN-Mot, DMN-VI, DMN-SC, Mot-Mot, Mot-VI, VI-SAL, VII-VII, VAs-SAL, SAL-SAL, SAL-SC, SC-CBL |
| FC/Change/Females | FC/Change/ Males | Females/ Males | positive | MF-MF, MF-DMN, MF-Mot, MF-SAL, MF-SC, FP-DMN, FP-Mot, DMN-Mot, DMN-CBL, Mot-CBL | MF-FP, MF-DMN, MF-CBL, FP-Mot, FP-SAL, FP-SC, DMN-SC, VII-SAL | MF-MF, MF-Mot, MF-SAL, MF-SC, FP-VI, DMN-VI, Mot-VII, Mot-VAs, Mot-SAL, Mot-SC, Mot-CBL, VAs-SC, VAs-CBL, SAL-SC |
| SC/T1/Females | SC/T1/Males | Females/ Males | negative | MF-FP, MF-Mot, FP-SC | FP-FP | MF-MF, MF-FP, MF-DMN, MF-Mot, MF-VI, MF-SAL, MF-SC, FP-DMN, FP-Mot, FP-VAs, FP-SC, FP-CBL, Mot-Mot, Mot-SAL, Mot-SC, VI-CBL, VII-SAL, VII-CBL, SAL-SAL, SAL-CBL, CBL-CBL |

*Note*: Subnetwork pairs with shared versus unique edges across paired CPM models. Each comparison contrasts two models. The two models being compared are represented as A and B, which includes functional connectivity (FC) vs structural connectivity (SC), cross-sectional or Time 1 (T1) vs prospective change score (Change), and males vs females. Shared edges score reflects the total magnitude (i.e., summed count) of edges common to both models. Shared pairs are the subnetwork pairs common to both models; *unique A* and *unique B* pairs are those specific to each model. Subnetwork pairs are reported separately for positive and negative networks. Only comparisons reaching significance in the main CPM analysis are shown. Abbreviations: MF = medial frontal; FP = frontoparietal; DMN = default mode network; Mot = sensorimotor; VI = visual A; VII = visual B; Vas = visual association network; SAL = salience; SC = subcortical; CBL = cerebellum/brain stem

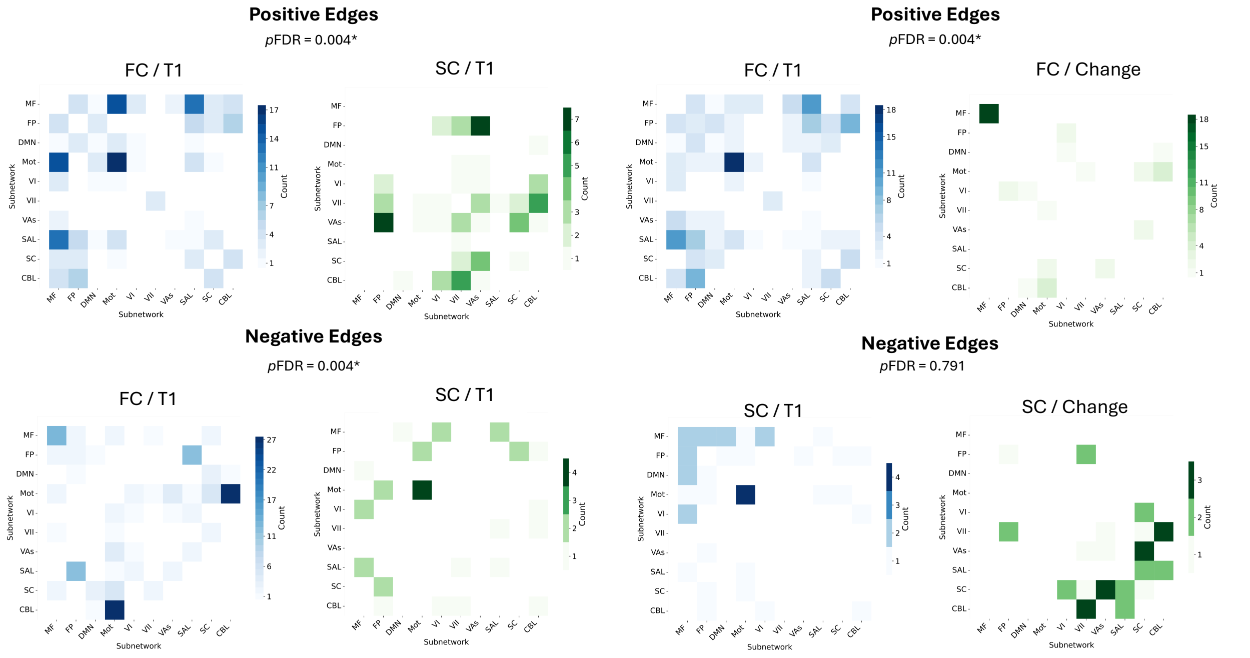


**Figure S1**. Unique network pairs for comparisons within the male subgroup. Network pairs are shown that are unique in each model comparison for males. The models being compared include functional connectivity (FC) vs structural connectivity (SC) and cross-sectional or Time 1 (T1) vs prospective change score (Change). Network pairs are reported separately for positive and negative networks. Only comparisons reaching significance in the main CPM analyses are shown. The $p_{FDR}$ values correspond to Supplementary Results Table S2.

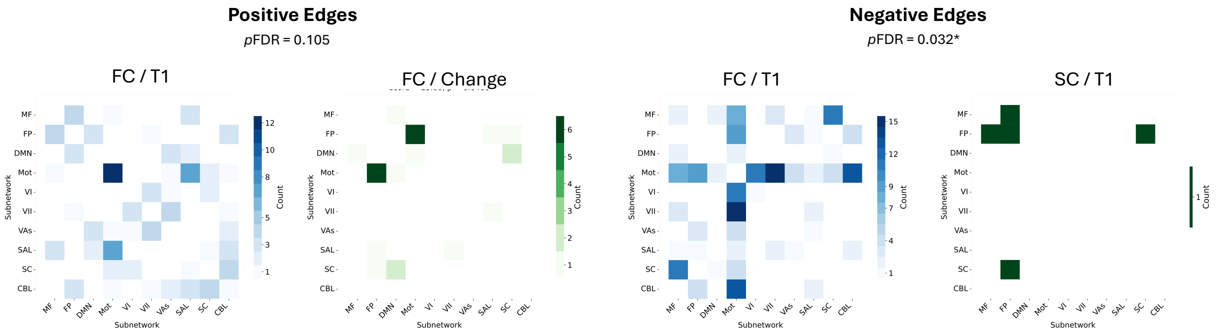


**Figure S2.** Unique network pairs for comparisons within the female subgroup. Network pairs are shown that are unique in each model comparison for females. The models being compared include functional connectivity (FC) vs structural connectivity (SC) and cross-sectional or Time 1 (T1) vs prospective change score (Change). Only comparisons reaching significance in the main CPM analyses are shown. The $p_{FDR}$ values correspond to Supplementary Results Table S2.

**Supplemental Discussion:**

**Sensorimotor Networks and Disruptive Behavior**

Brain-behavior associations were also observed in the sensorimotor and cerebellar networks. The sensorimotor network contributes to the integration of sensory and motor information supporting goal-directed actions, movement, and processing of bodily states [65]. Alterations in the sensorimotor network has also been linked to psychopathology more broadly [66]. These findings are consistent with our work implicating sensorimotor networks implicated in the prediction of childhood disruptive behavior during emotional face processing [34] and with prior work linking sensorimotor connectivity to irritability/anger during frustration [67]. The present findings extend prior work by showing that sensorimotor connectivity contributed to concurrent and prospective models through connections with cerebellar, medial frontal, frontoparietal, and visual networks. Thus, disruptive behavior may be associated with distributed connectivity among higher-order control systems and systems supporting sensory, motor, and affective responding. That is, connectome dysfunction may progress from local disruptions prior to seemingly disparate networks that are globally connected to frontoparietal emotion regulation circuitry [68], which could suggest a 'cascading network failure' theory implicated in disruptive behavior [69,70].

**Sex-Specific Connectome Markers of Disruptive Behavior**

Sex differences in connectome markers were observed in prediction performance, modality, and network patterns. Functional connectivity predicted disruptive behavior severity in both females and males for concurrent and prospective models, whereas structural connectivity predicted disruptive behavior in males but not in females. Effect sizes for functional connectivity also tended to be higher than structural connectivity. While recent studies have not reported sex differences in neuroanatomical markers of disruptive behavior in youth [14,15], several studies have observed sex-by-diagnosis interactions in brain structure [18-21,23] and functional connectivity [24-27]. In our work, we showed that girls and boys with disruptive behavior problems show distinct patterns of cortical volume and thickness in frontoparietal nodes and frontolimbic functional connectivity in clinically affected youth with disruptive behavior problems relative to unaffected controls [19,26]. Our findings are also consistent with prior work indicating sex differences in networks linked to disruptive behavior [28], our findings suggest sex-specific patterns in disruptive behavior as well as shared patterns across females and males. For instance, the default mode contributed predicted edges across models and in females and males. Additionally, while our functional connectivity findings are in line with prior work showing that disruptive behavior can be predicted across the sexes [17], our findings here suggest a more nuanced pattern of sex-specific connectome markers. For example, within and between network connectivity with medial frontal network showed a distributed positive connectivity pattern predicting concurrent disruptive behavior severity, but a more focal positive connectivity pattern predicting worsening disruptive behavior in the total and male subgroup compared to the female subgroup. In contrast, the female subgroup showed more focal pattern of negative connectivity between sensorimotor-visual networks cross-sectionally and positive connectivity between the frontoparietal and sensorimotor networks. Sex differences in structural connectomes were also observed. Predictive models were significant for males but not females. This finding is also consistent with prior work reporting associations between abnormal microstructural integrity and disruptive behavior (anger) in males but not females [52]. In the present study, predictive models in male and female subgroups were balanced and relatively greater severity of disruptive behavior in males vs females could impact predictive models training and performance. Thus, future work in a sex-balanced, clinically affected sample of youth will be important to provide a range of severity extending into clinically significant range to test sex-specific markers of disruptive behavior.

**Multimodal and Model-Specific Connectome Organization**

Supplemental follow-up comparisons of predictive feature masks indicated partially distinct connectome configurations across functional vs structural modalities and across concurrent vs prospective models. While no shared connectivity comparisons survived FDR correction, several model-specific contributions remained significant. Thus, these findings suggest that functional and structural connectivity may provide complementary insights about disruptive behavior rather than converging on a single predictive connectome. The observed limited edge overlap is plausible given that functional and structural connectivity reflect related but distinct aspects of neural architecture: functional connectivity captures coordinated activity across distributed nodes, whereas structural connectivity reflects microstructural properties of reconstructed anatomical tracts. The two neuroimaging modalities may therefore capture distinct, non-overlapping predictive connections across circuit systems underlying emotion regulation, broader cognitive control, sensorimotor, and social perception. Future work integrating functional and structural features within a fused or unified multimodal model could test whether complementary contributions improve prediction beyond either modality alone.

**Connectome Markers of Concurrent vs Prospective Disruptive Behavior**

Predictive performance across imaging modalities tended to be higher for the concurrent than prospective models. Additionally, predictive edges for concurrent models were more widely distributed, while prospective models showed a more focal pattern. For functional and structural connectomes, medial frontal and frontoparietal networks contributed to concurrent and prospective prediction, but differences emerged in the extent to which these nodes contributed across time points and modalities, and coordination across other networks including salience, default mode, cerebellar, and sensorimotor circuits. These findings could suggest that partially distinct patterns of circuit markers predict current symptom severity and subsequent symptom change, which is consistent with prior studies reporting different neural markers of concurrent and prospective disruptive behavior symptoms [93,94]. Prior longitudinal neuroanatomical studies of disruptive behavior have suggested attenuated development of lateral prefrontal, frontoparietal, and frontolimbic regions implicated in disruptive behavior [53,93,95]. Consistent with neuroanatomical work, in the current study structural connectivity findings were driven largely by predictive negative edges, indicating reduced baseline white matter integrity was associated with greater symptom severity and worsening symptoms, which may suggest attenuated integrity of networks implicated in disruptive behavior problems [50-53,95]. However, the present study used baseline connectomes rather than repeated imaging and therefore cannot determine whether altered developmental changes in connectivity were associated with symptom changes. Future studies using a repeated neuroimaging design in a clinical sample of youth will be necessary to distinguish stable circuit markers of risk and vulnerability from connectivity changes associated with persisting, increasing, or desisting symptom courses.
